# Limits of deep-learning-based RNA prediction methods

**DOI:** 10.1101/2025.04.30.651414

**Authors:** Marko Ludaic, Arne Elofsson

**Affiliations:** Science for Life Laboratory and Department of Biochemistry and Biophysics, Stockholm University, Solna 171 21, Sweden

**Keywords:** RNA structure, Deep Learning, Structure Prediction, RNA-Protein interactions

## Abstract

In recent years, tremendous advances have been made in predicting protein structures and protein-protein interactions. However, progress in predicting RNA structure, either alone or in complex with other macromolecules, has been less prominent, though some recent developments have been reported. It remains unclear whether the improved prediction accuracy is sustained for novel RNA structures. Here, we use an independent benchmark to evaluate the performance of the latest methods. First, we show that state-of-the-art methods can sometimes predict the structure of single-chain RNA strands, with accurate models observed for RNAs with well-defined or regular secondary structures. Next, our evaluation was extended to RNA complexes, where prediction accuracy was notably higher for those involving extensive canonical base pairing. Additionally, a structural similarity analysis revealed that prediction success strongly correlates with resemblance to known structures, indicating that current methods recognise recurring motifs rather than generalising to novel folds. Finally, we also noted that the accuracy estimates for RNA models are far from accurate. Therefore, it is not possible to reliably identify the correctly predicted models with today’s methods.

**Graphical Abstract:** 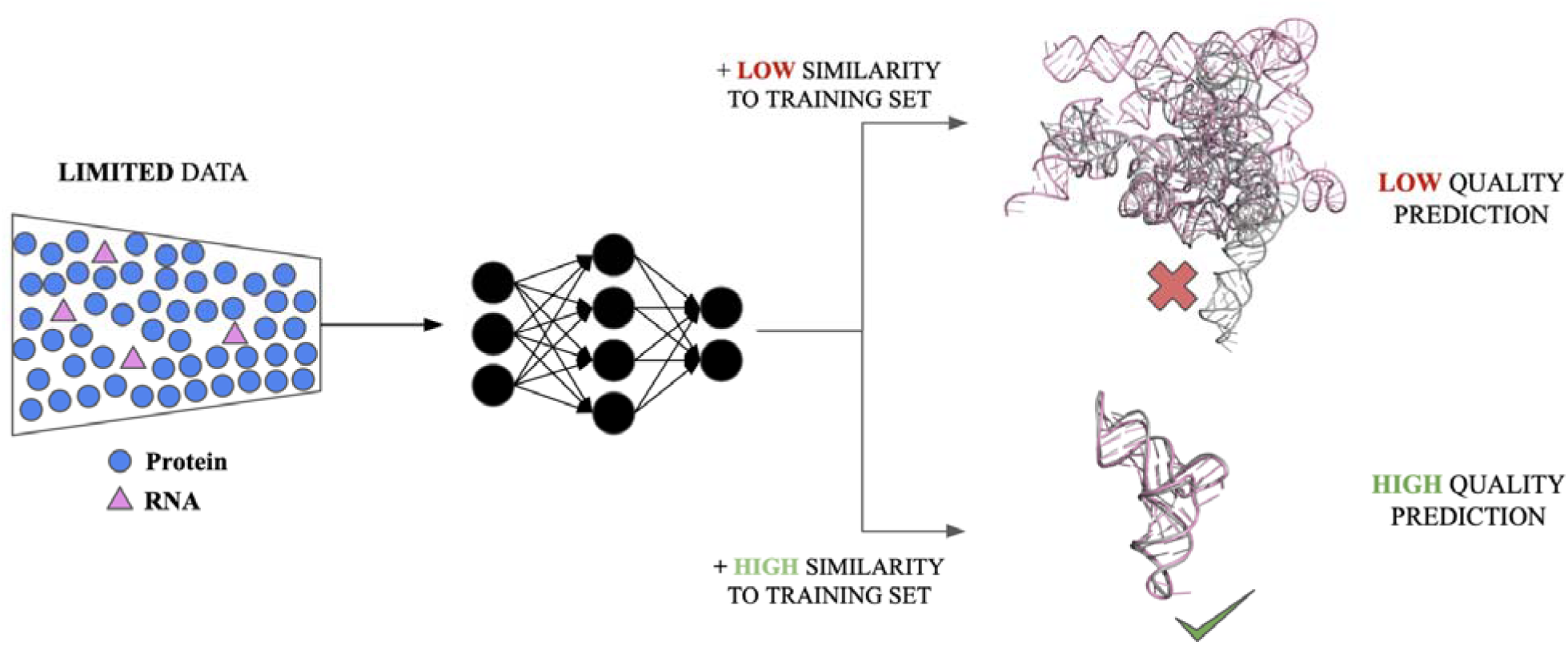

## Introduction

RNA molecules are highly dynamic macromolecules that adopt intricate three-dimensional (3D) structures essential for many cellular and viral processes, emergent RNA-based medicines, and biotechnologies. Different types of RNA, such as antisense oligonucleotides, small interfering RNAs, and microRNAs, have shown promising therapeutic potential (1). Even though RNAs were often viewed as mere intermediaries in the transfer of genetic information from DNA to protein synthesis, they are also recognised for their functional versatility, extending far beyond their linear nucleotide sequences.

RNA structures are critical for elucidating structure-function relationships; therefore, modelling RNA structure from its nucleotide sequences remains an important and largely unsolved task. There are two main reasons for the limited success. First, the limited availability of determined RNA structures, at least one order of magnitude less than for proteins, poses a significant challenge for training AI-based models. Secondly, RNA’s conformational dynamicity complicates the application of deep learning models. Many RNAs undergo drastic conformational changes to obtain a shape required for binding to a molecular partner (2). This is also one of the reasons why experimental methods such as X-ray crystallography, NMR, and cryo-EM have primarily been able to resolve regulatory and enzymatic RNAs such as tRNA, rRNA, and ribozymes (3). The structure of other RNA types, such as long non-coding RNAs (lncRNAs), remains less well studied, possibly because they are indeed more disordered.

Before the release of RoseTTAFold2NA (RF2NA) in early 2024 (4) and AlphaFold3 later in 2024 (5), deep learning approaches were limited to predicting the structures of monomeric RNAs and could not handle RNA in a complex with other molecules. Recently, several deep learning approaches based on the AlphaFold3 architecture, such as Boltz-1 (6), Chai-1 (7), and HelixFold3 (8), were released. These methods can handle proteins, nucleic acids, small molecules, and ions, greatly expanding the scope and applicability of structure prediction. Furthermore, some methods for predicting single-chain RNA, such as RhoFold+ (Shen et al., 2024), utilize large language models (LLMs) to predict atom coordinates, while others are adaptations of AlphaFold. Table 1 provides an overview of the latest methods for predicting RNA structure.

**Table 1.** A brief description of the most critical characteristics of benchmarked methods for single-chain RNA and RNA-RNA/Protein prediction.

| METHOD | DESCRIPTION | RNA |
| --- | --- | --- |
| AlphaFold 3 | A deep learning method consisting of a pairformer for residue interaction modeling and a diffusion module for denoising raw atomic coordinates, removing global rotational/translation constraints to enhance prediction flexibility and accuracy. | Single, complex |
| Boltz-1 | AlphaFold 3-inspired architecture with modified order of operations in the MSAModule and transformer layer. | Single, complex |
| Chai-1 | AlphaFold 3-inspired architecture incorporating residue-level embeddings from a large language model for proteins, also introducing experimental constraint features like pocket, contact, and docking constraints. Trained without RNA MSAs. | Single, complex |
| HelixFold3 | AlphaFold 3-inspired architecture using PaddlePaddle library allowing for faster model training. | Single, complex |
| NuFold | A deep learning method built upon AlphaFold architecture design to predict RNA structures by incorporating nucleic acid sequences, secondary structure inputs, and RNA-specific inter-base angles, distances, and atom positions. | Single |
| RhoFold+ | A deep learning model for RNA structure prediction that utilizes an RNA language model, alongside several deep learning modules, including an IPA module for modeling 3D positions. | Single |
| RoseTTAFoldNA | A three-track architected model that simultaneously updates sequence (1D), residue-pair (2D), and structural (3D) representations to predict macromolecular structures.. | Single, complex |
| trRosettaRNA | A deep learning model consisting of MSA and pair representations fed into a transformer network (RNAformer) predicting 1D and 2D geometries, refined in the final 3D structure folding step. | Single |

Until recently, systematic benchmark studies directly comparing state-of-the-art methods have been scarce. Nevertheless, community-wide assessments such as RNA-Puzzles Round V (9) and the CASP16 (Critical Assessment of Structure Prediction) RNA assessment (10) have been conducted. These efforts are organized as competitions in which participating groups submit predictions for blind targets where the focus is on evaluating the performance of the participants, rather than on a systematic comparison of available methods under identical input conditions. To our knowledge, one study (11) compared RNA structure prediction methods for single-chain RNA. However, it did not include many of the most recent approaches (e.g., Boltz-1, HelixFold 3, Chai, RosettaFoldNA, and NuFold). Additionally, another recent study benchmarked RNA–RNA interaction methods limited to secondary structure prediction (12), focusing on base-pair prediction accuracy rather than three-dimensional structure modeling. The most recent benchmark also included RNA-protein complexes, but compared only two methods, AlphaFold3 and RosettaFoldNA (13).

## Materials and Methods

### Benchmark dataset

RNA-containing entries released after September 30, 2021 (for RNA complexes) and February 28, 2022 (for single-chain RNAs), which correspond to the latest training set date cutoff of the benchmarked methods, were downloaded from the Protein Data Bank (PDB) along with their corresponding sequences in January 2026. For each entry, the first biological unit was used. In addition, we used CASP16 targets containing RNA to supplement the benchmark dataset.

All sequences were filtered using CD-HIT-EST (14) at 80% sequence identity to ensure dataset diversity. The clustering was performed both within the benchmark dataset and against the remaining RNA sequences in the PDB deposited before the training date cutoff. Sequence identity was calculated using the longer sequence as a reference, which was critical, as many RNA sequences in the PDB are synthesized from short RNAs. Applying this criterion avoids falsely reporting 100% sequence identity between a short and a long RNA sequence. Sequences containing ambiguous nucleotides, such as’X’ or’N’, were removed from the dataset. Furthermore, sequences shorter than 10 nucleotides that consisted predominantly of a single nucleotide type (>80% homogeneity) were also excluded, as they are likely to be linear and lack a modular structure, making them unsuitable for inclusion in the benchmark dataset.

These filtering criteria were applied to both single-chain RNAs and RNA sequences from RNA complexes.

To evaluate structural redundancy, RNA-align TM-score (TM-scoreRNA) (15), a length-dependent variant of TM-score optimized for RNA molecules, was used. Structures of RNA complexes were divided into individual chains, RNA chains extracted and processed together with single-chain RNA structures. Since there is no RNA-specific tool for structural similarity search equivalent to MMseqs2 (16) for proteins, the pairwise TM-score was calculated between all structures within the initial benchmark dataset, and those with a TM-score > 0.7 were considered structurally too similar and excluded from the benchmark dataset. Each case was also manually reviewed to confirm that TM-score values accurately reflect actual structural similarity. TM-scores were also calculated against the training set. This procedure finds structurally similar folds that models have seen during the training. We preserve those structures and discuss how dependent the model’s accuracy is on the training set in the last section of results (See *Results and Discussion, Prediction accuracy dependence on AlphaFold3 training set*). In the case of RNA-protein complexes, protein structural similarity was not used as a filtering criterion, as doing so would reduce the dataset size. Instead, the filtering remained focused on RNA, consistent with the benchmark’s aims of evaluating RNA prediction accuracy, rather than protein structure prediction.

Finally, the dataset was further reduced because several sequences in the initial test set could not be modelled by specific methods. Modelling complexes with more than four protein and/or RNA subunits with Boltz-1 version 0.4.1 failed due to out-of-memory errors. Consequently, such complexes were excluded from the analysis, reducing the benchmark dataset for complexes to 158 structures. After filtering, our benchmark dataset for single-chain RNA structures comprised 86 models.

### Model generation

The benchmark evaluates the performance of eight prediction methods, AlphaFold3 (5), Boltz-1 (6), Chai-1 (7), HelixFold3 (8), NuFold (17), RhoFold+ (18), RoseTTAFoldNA (4), and trRosettaRNA (19) for single-chain RNA. Four of the methods. AlphaFold3, Boltz-1, HelixFold3, and RoseTTAFoldNA also predicted RNA-RNA and RNA-protein complexes. Chai-1, a multimodal foundation model for molecular structure prediction, was excluded from the RNA complex analysis due to its 2048-token inference limit. We also initially included DRFold (20), however, it consistently failed to generate structures exceeding 205 nucleotides in length. Excluding those structures would have substantially reduced the representativeness of our benchmark data. Therefore, we excluded DRFold from the analysis. The same was done with DeepFoldRNA (21), which failed to generate full-atom models, preventing proper evaluation of its predictions.

For each method, the structures were generated on a single NVIDIA DGX-A100 Core GPU using the latest version (v1.0 for trRosettaRNA) as of October 1, 2025, with default settings. For all methods, the same set of MSAs was used. These were generated using rMSA (22) for RNA sequences and MMseqs2 (16) for protein sequences. An rMSA search was performed against the RNACentral and NT databases using default settings. For proteins, MMseqs2 was run on the ColabFold databases as of October 1, 2025, which consist of UniRef30, BFD/Mgnify, and ColabFold DB (23). MSAs were converted into alternative formats depending on each model’s specific requirements. The conversion scripts are available from the accompanying git repository. For each method, five models with the same seed were generated for each target.

The top-ranked model was used for analyzing each method. Several methods, including AlphaFold3, report the predicted template modeling (pTM) and the interface predicted template modeling (ipTM) scores in the output, and internally rank models. Other methods, such as RF2NA or RhoFold+, report the predicted local distance difference test (pLDDT), which reflects confidence in the local structure (24). We ranked the models based on the average pLDDT scores for these methods, as this was the score that was produced by the network. If a method did not report any confidence or ranking score, we selected the first structure. This approach aligns with the practical goal that the top model in the output should ideally represent the best prediction. We also noted that, in almost all cases, the quality difference between generated models is minimal (see Figures S1-S4).

### Model evaluation

To evaluate the quality of predicted models, we used four different scores that were also used in the latest CASP16 assessment of three-dimensional RNA structure (10). More specifically, for single-chain RNAs, we calculated TM-score via RNA-align (15), Interaction Network Fidelity (INF), Local Distance Difference Test (lDDT) and a Global Distance Test (GDT-TS). To calculate INF, lDDT, and GDT-TS, we used RNAdvisor (24), a comprehensive benchmarking tool for measuring and predicting RNA structural model quality. For assessing RNA complexes, in addition to the US-align TM-score, we used the DockQ score (25), which was also used in the previously mentioned CASP evaluation of nucleic acids. DockQ provides an overall score, averaged across all interfaces, as well as individual scores, one for each interface. We also assessed the accuracy of RNA-RNA and RNA-protein interfaces within each complex, using the per-interface DockQ score. In PDB entries, the deposited sequence (provided in FASTA/SEQRES records) does not always fully correspond to the experimentally resolved structure, as flexible or disordered regions may be missing in the atomic coordinates. As a result, predicted models and reference structures may differ in length or residue composition. To ensure a consistent and fair comparison, predicted structures were trimmed to match the resolved residues present in the reference structure prior to evaluation.

We calculate the TM-score using the following command line options:

`RNAalign {model.pdb} {reference.pdb}`

for single-chain RNA molecules and

`USalign {model.pdb} {reference.pdb}-mm 1-ter 0`

for RNA complexes.

## Results and Discussion

### Performance of structure prediction methods on single-chain RNAs

Figure 1 shows the performance of deep learning methods for single-chain RNA prediction. Among the tested methods for single-chain RNA prediction, AlphaFold 3 was a top performer based on INF (0.709) and lDDT (0.6), while Boltz-1 was the best model based on GDT-TS (0.481). Overall, both AF3 and Boltz-1 achieved a mean TM-score of 0.326 and a success rate of 19% and 14%, respectively. This is followed by NuFold (0.325/19%), RF2NA (0.296/10%), Chai (0.291/12%), HF3 (0.287/8%) and RhoFold+ (0.276/12%). (Figure 1A). An RNA model is considered successful if its TM-score is above 0.45; the same criteria were used to assess nucleic acid structure prediction in CASP16 (10). Scores for all models are shown in Tables S1-S4, while all mean scores and success rates are shown in Table S5. To better understand performance differences across RNA types, we plotted the TM-scores of the top-performing method, AlphaFold3, against those of all other methods. A paired T-test reported no significant difference between the predictions from AF3 and Boltz-1 (Figure 1B; Cc = 0.85, P = 9.97*10-1), but HF3 (P = 8.77*10-4), RF2NA (P = 1.71*10-2), RhoFold+ (P = 1.76*10-7), trRosettaRNA (P = 1.77*10-6) and Chai (P = 7.22*10-4) are statistically significantly less accurate than AF3 (Figure S5).

**Figure 1.**
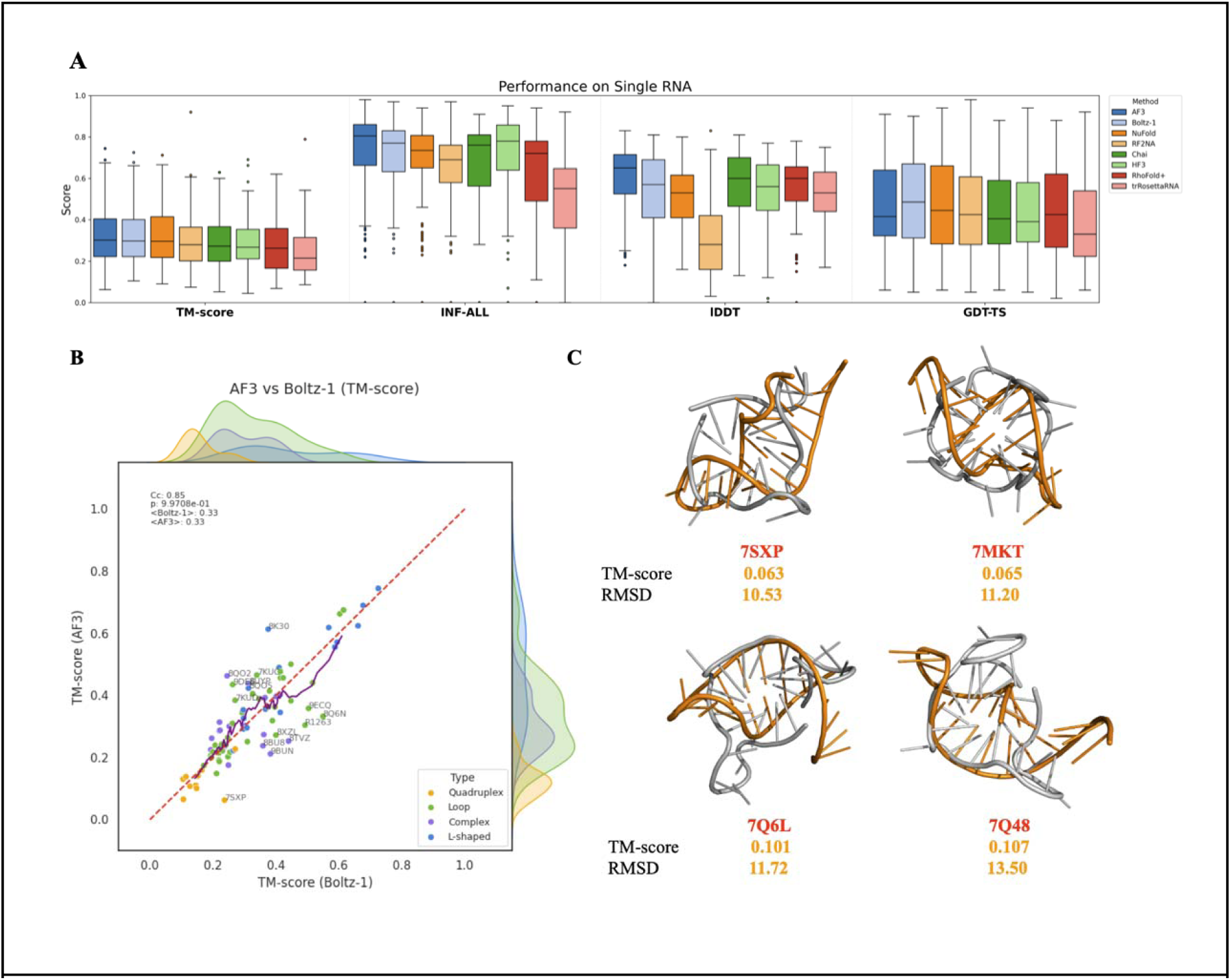
Performance comparison for different RNA prediction methods on single-chain RNA. (A) The performance of RNA prediction models was evaluated using four different evaluation metrics (TM-score, INF-ALL, lDDT, GDT-TS). The distribution for each score and method is presented as a box plot, with the median marked by a line and the interquartile range shown. The methods are arranged in descending order of the mean TM-score. Different colors correspond to the various methods. (B) Pairplot showing correlation between Boltz-1 (x-axis) and AlphaFold3 (y-axis) TM-scores for single-chain RNA. The labeled points show PDB IDs of predicted models whose difference in TM-score between the two methods is greater than 10%. Paired t-test reported no significant difference between the two methods (Cc = 0.85, P = 9.97*10^-1^). (C) Visualization of the four least accurate AF3 predictions. Ground-truth structures are colored in grey, while the predicted structures are shown in orange.

When RNAs were grouped into four structural classes: short RNA loops, G-quadruplexes, L-shaped conformations such as tRNAs, and complex RNAs composed of multiple secondary-structure motifs, we noticed apparent differences in prediction accuracy (Figure 1B). In general, RNAs with well-defined and relatively simple folds were predicted more accurately than structurally complex RNAs. Structures that adopt common folds that are well represented in PDB, such as L-shaped RNAs, were modeled most accurately across all methods (Figure S5). In contrast, small G-quadruplex RNAs consistently ranked among the worst-scoring targets. The four lowest-scoring single-chain RNAs generated using AlphaFold 3 are shown in Figure 1C. Observing the predicted models superposed to ground truth structures, it is noticeable that AF3 recognizes the quadruplex fold, but fails to model it correctly. In addition, one G-quadruplex RNA, 7SXP, was modeled slightly less inaccurately by Boltz-1 (TM-score = 0.237) than by AF3 (TM-score = 0.063). The structure 7SXP was released in May 2023 and was not a part of the training set for either method. Although Boltz-1 achieved higher GDT-TS (0.79) and INF (0.42) values for 7SXP, the overall difference in model quality is moderate, and TM-scores remain low for both models (Figure S6). This behaviour is consistent with the limitation of TM-score when applied to short RNAs and G-quadruplex structures.

### Length-Dependent Limitations of TM-score in RNA Structure Evaluation

To examine success rates, we defined thresholds for each metric in the same way as was done in th CASP16 evaluation of nucleic acids (10). When examining success rates using thresholds of TM-score > 0.45, GDT-TS > 0.45, INF > 0.75 and lDDT > 0.75, we observed that among the two global fold metrics (GDT-TS and TM-score), GDT-TS consistently yields substantially higher success rates than TM-score across all methods (Figure 2A). In other words, many models that are classified as correct by GDT-TS are not considered successful by TM-score. To better understand this discrepancy, we examined structures with high GDT-TS but low TM-score. The four most pronounced examples are shown in Figure 2B. These structures also exhibit high INF values, indicating well-captured base-pairing, stacking and local interactions. Therefore, low TM-scores are not necessarily reflective of a poor structural quality, but rather a limitation of the TM-score when applied to short RNA chains. This behaviour comes from the normalization by the length-dependent scaling factor (dlJ) used in the TM-score formulation (15). For short RNAs (e.g., < 40 nt), the scaling factor becomes so small that even minor structural deviations lead to disproportionately low TM-scores. Also, lDDT is much more restrictive than INF among the local measures, likely due to an lDDT threshold of 0.75 being quite strict, in particular for models generated by some methods (trRosettaRNA, RF2NA).

**Figure 2.**
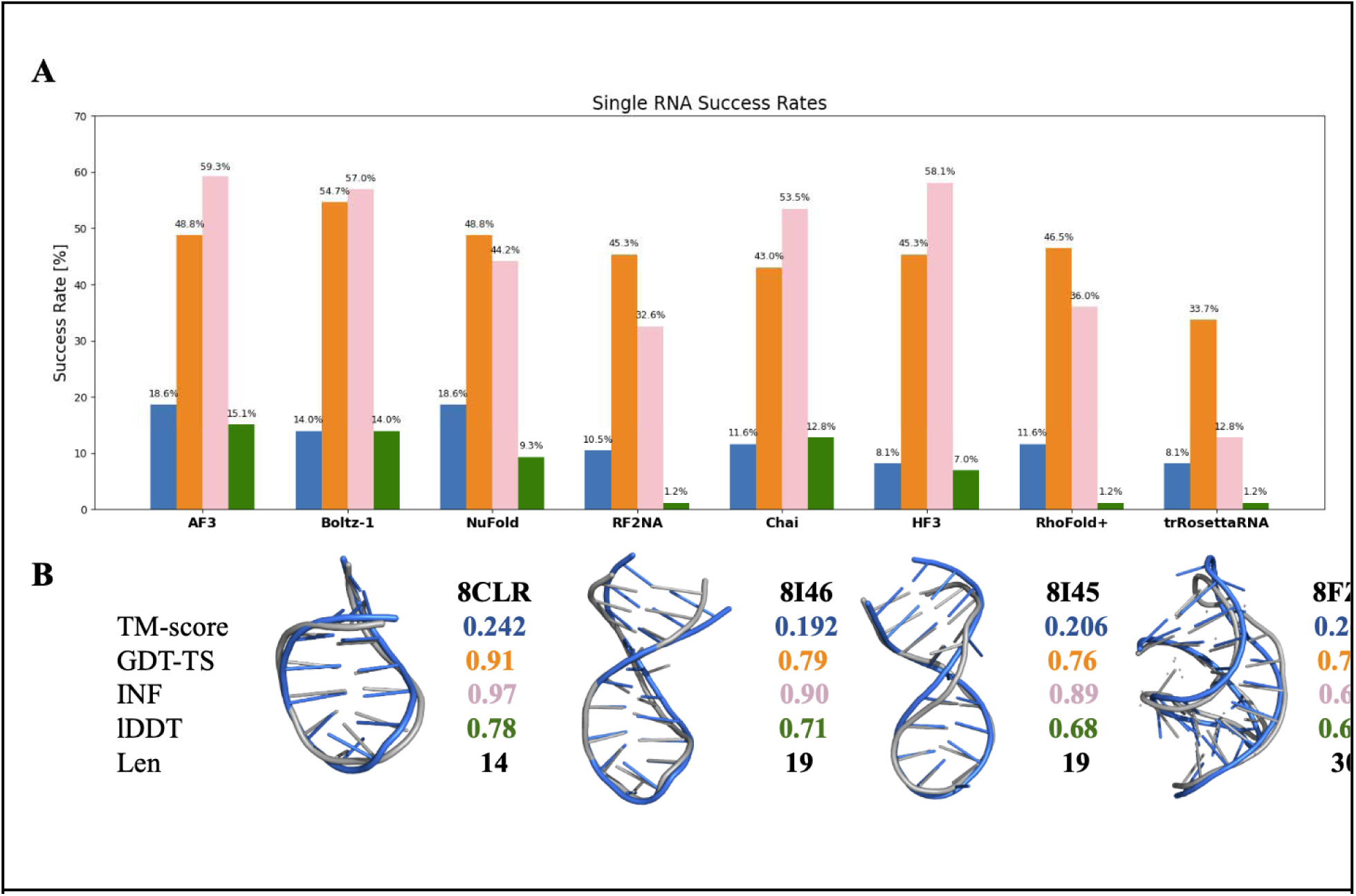
(A) Success rates achieved by all methods are defined as the fraction of acceptable models with a minimum GDT-TS and TM score of 0.45, and lDDT and INF score of 0.75. (B) Four models with a higher GDT-TS and INF, but very low TM-score. The ground-truth structures are colored in grey, while predicted structures are shown in blue.

A previous RNA benchmarking study (26) also noted this issue and addressed it by excluding all RNAs shorter than 60 nucleotides from their benchmark. However, we consider this filtering undesirable for several reasons. Many biologically important RNAs are short yet structured, and excluding them removes a meaningful part of RNA structural space. Moreover, because the methods evaluated in this benchmark were trained on the entire PDB, which includes numerous RNAs shorter than 60 nt, excluding short RNAs from the benchmark would prevent a fair assessment of model performance on sequences that were part of their training distribution.

Despite the limitations imposed by length constraints, we retain TM-score in our analysis. This choice is motivated by its widespread use, its RNA-specific optimization, and its ability to capture global structural similarity in a consistent manner across methods. Nonetheless, we note that the length dependence of TM-score becomes unreliable for short RNAs, and the results in this range must be used carefully. Instead of discarding short RNAs, we adopt a more robust definition of a successful prediction: we consider a prediction successful if any individual model exceeds the predefined success thresholds for each metric. We then analyze the overlap between these criteria using Venn diagrams, quantifying how many structures are consistently classified as successful across multiple metrics. When aggregating all predictions across all methods, only 12 models satisfy the success thresholds for all four metrics simultaneously (Figure 3A). Out of the 12 most successful predictions, the top 3 predictions shown in Figure 3B include one tRNA-like structure (R1263) and two helical RNA structures (8FCS modelled by AF3 and 8FCS modelled by NuFold). In contrast, 71 models pass a threshold on at least one metric, and 42 models are classified as correct using both INF and GDT-TS, suggesting that individual metrics alone may overestimate accuracy. These results underscore the importance of using multiple evaluation metrics for reliable assessment of RNA structure prediction quality. Venn diagrams for individual methods are given in the supplementary material (Figure S7-8).

**Figure 3.**
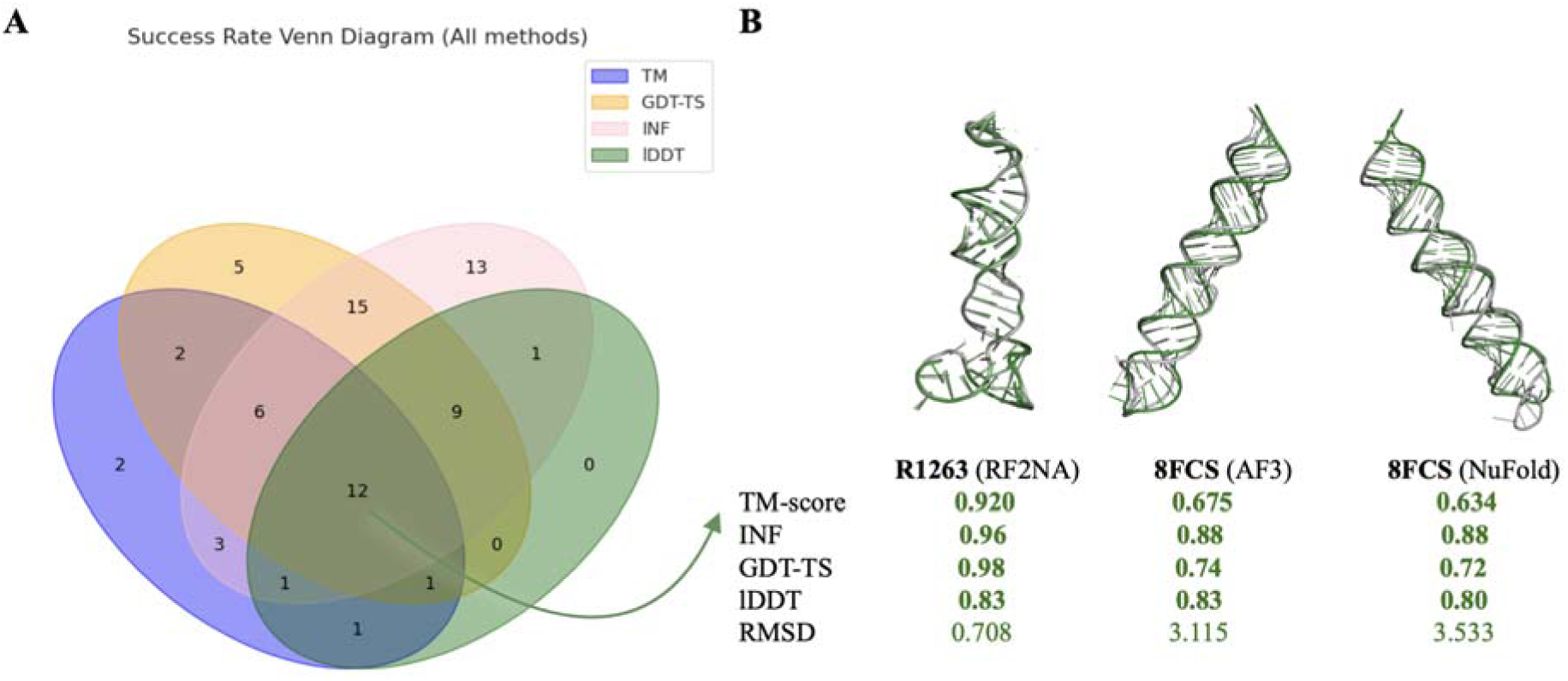
(A) Venn diagram showing the number of models passing the thresholds for different evaluation metrics, for all predictions across all methods. Each color represents one metric. (B) Visualization of the top three predicted structures (8VT5, 8FCS, 9BLM), selected based on the defined success thresholds.

To further investigate the influence of RNA length on structure accuracy, we analyzed pairwise relationships among the four evaluation metrics. Figure 4 shows all six pairwise correlations with data points colored by RNA length. The dataset is divided into three length intervals: short RNAs (13-28 nt), medium RNAs (29-68 nt), and long RNAs (69-497 nt). Dark blue points correspond to short RNA structures, lighter blue medium-sized RNAs and red points to the long RNAs. Grey dashed lines indicate the thresholds used to define the success rate for each metric, as previously defined.

**Figure 4.**
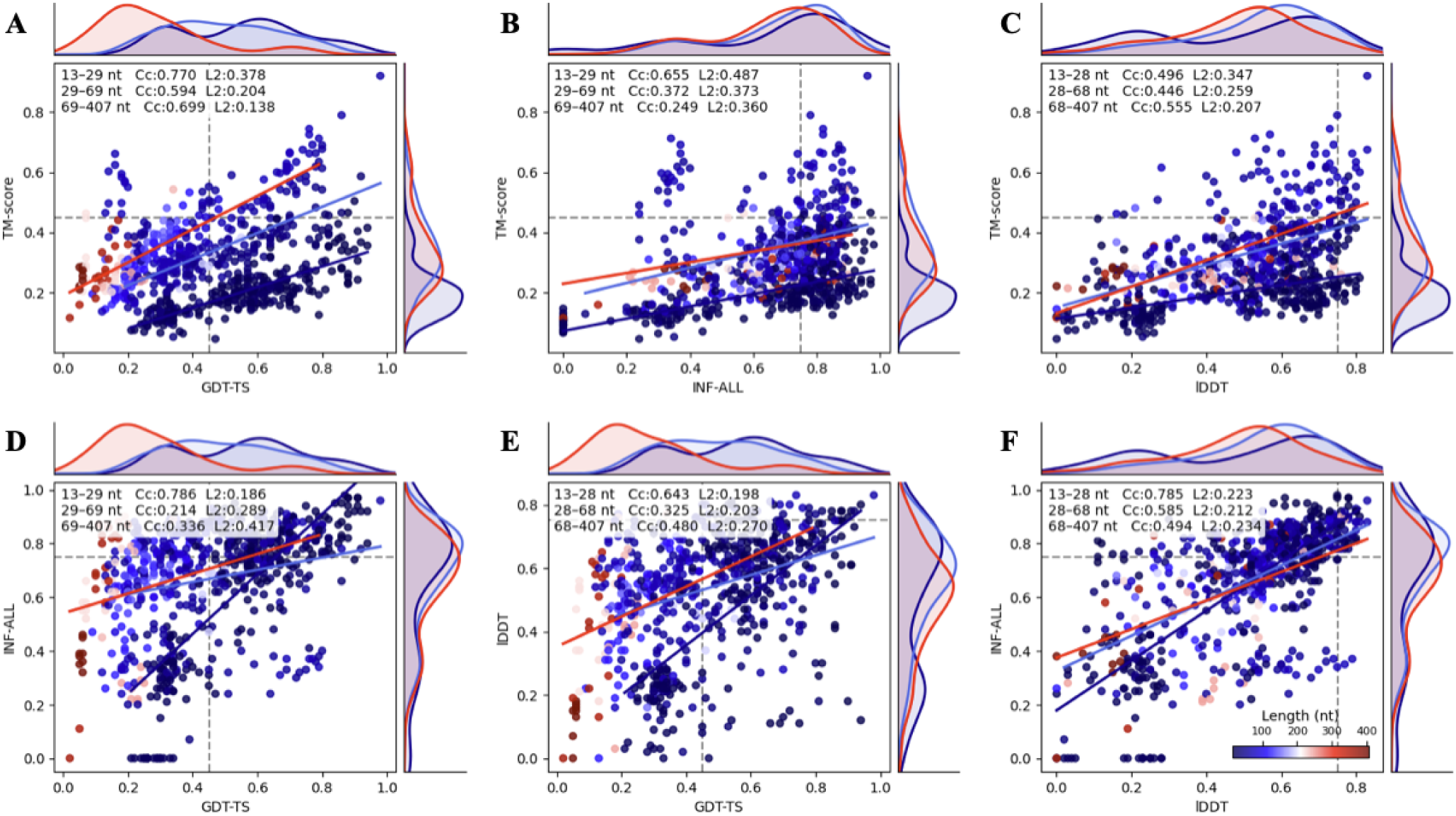
Pairwise comparison of evaluation metrics. (A) Correlation between TM-score and GDT-TS. (B) Correlation between TM-score and lDDT (C) Correlation between TM-score and INF. (D) Correlation between INF and GDT-TS (E) Correlation between lDDT and GDT-TS. (F) Correlation between INF and lDDT. Each plot is divided into three length intervals. Cc denotes the Pearson correlation coefficient with L2 error also reported.

Comparing correlations between TM-score and the other three metrics, a pronounced length-dependent effect is observed. Many short models fall into the lower-right quadrant, i.e., exceeding the GDT-TS (Figure 4A) and INF (Figure 4B) thresholds but being below the TM-score threshold. This indicates that short RNA models achieve relatively high GDT-TS and INF values despite low TM scores. In contrast, as RNA length increases towards medium-sized and long RNAs, the number of models in this quadrant decreases substantially. This trend is also noticeable in the distribution of TM-scores, which shifts towards higher values for longer RNAs. Consequently, TM-score shows stronger agreement with other evaluation metrics for longer RNAs, and models that exceed GDT-TS or INF threshold are more likely to also exceed the TM-score threshold.

Correlations between other pairs of metrics are also shown in Figure 4, with the strongest agreement observed between lDDT and INF (Figure 4F), both of which emphasize local structural accuracy. However, it can be seen that the threshold for lDDT is significantly stricter than for INF, as many models have INF-scores around 0.8 and lDDT scores around 0.6. This suggests that some models may still contain suboptimal local geometries. INF is the only metric entirely length independent, as observed from the INF score overlapping distributions for each length interval. On the other hand, GDT-TS shows a bimodal distribution for the shortest RNAs. Overall, this analysis highlights the importance of evaluating RNA structure predictions using multiple complementary metrics.

### Model quality dependence on MSA depth

Many deep-learning methods rely on MSAs to capture evolutionary relationships, detect distant homologs, and improve model accuracy. For all methods that incorporate MSA during inference, we computed MSA depth using NEFFy (27). Across all tested methods, we observe a positive correlation between MSA depth and model quality, as measured by TM-score. The greatest improvement was observed for AlphaFold3 and NuFold (Figure 5). However, for individual targets, the correlation remains moderate, suggesting that although deeper MSAs generally improve prediction accuracy, substantial gains are primarily observed for targets with very high sequence coverage.

**Figure 5.**
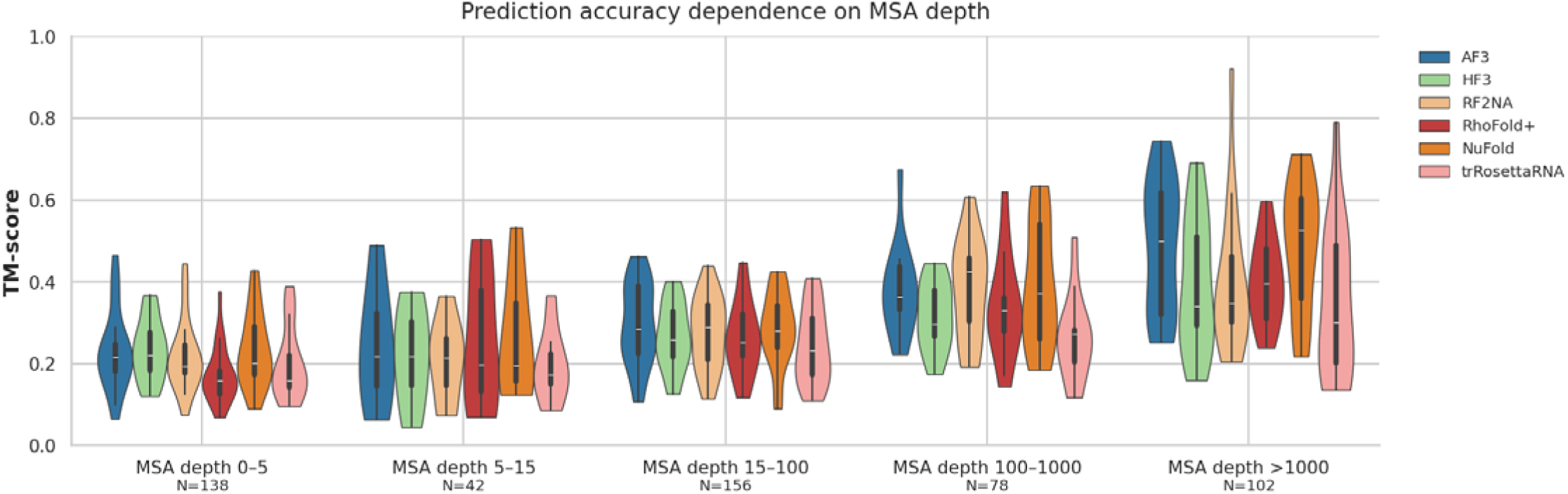
Prediction accuracy dependence on MSA depth. Distribution of TM-score across 5 MSA depth categories. *N* shows the total number of models in each category, across all methods. Results are shown for single-chain RNA prediction methods that incorporate MSAs during inference. Each color corresponds to a different method.

### Performance of structure prediction for RNA-RNA and RNA-protein complexes

The performance of the benchmarked methods, evaluated using TM-score and DockQ score, is summarized as violin plots in Figures 6A and 6B, respectively. Among the four methods, AlphaFold3 and Boltz-1 stand out, achieving mean TM-scores of 0.711 and 0.680, and mean DockQ scores of 0.390 and 0.357, respectively. In contrast, RF2NA and HF3 showed substantially lower performance, with mean TM-scores of 0.485 and 0.062, and mean DockQ scores of 0.391 and 0.166, respectively. Complete results for all benchmarked structures are provided in Tables S6 and S7. Using a DockQ score ≥ 0.23 to define successful models, the success rates for benchmarked methods are 54.4%, 54.4%, 12.0%, and 24.7% for AlphaFold3, Boltz-1, HF3, and RF2NA, respectively. All mean scores are shown in Table S8.

**Figure 6.**
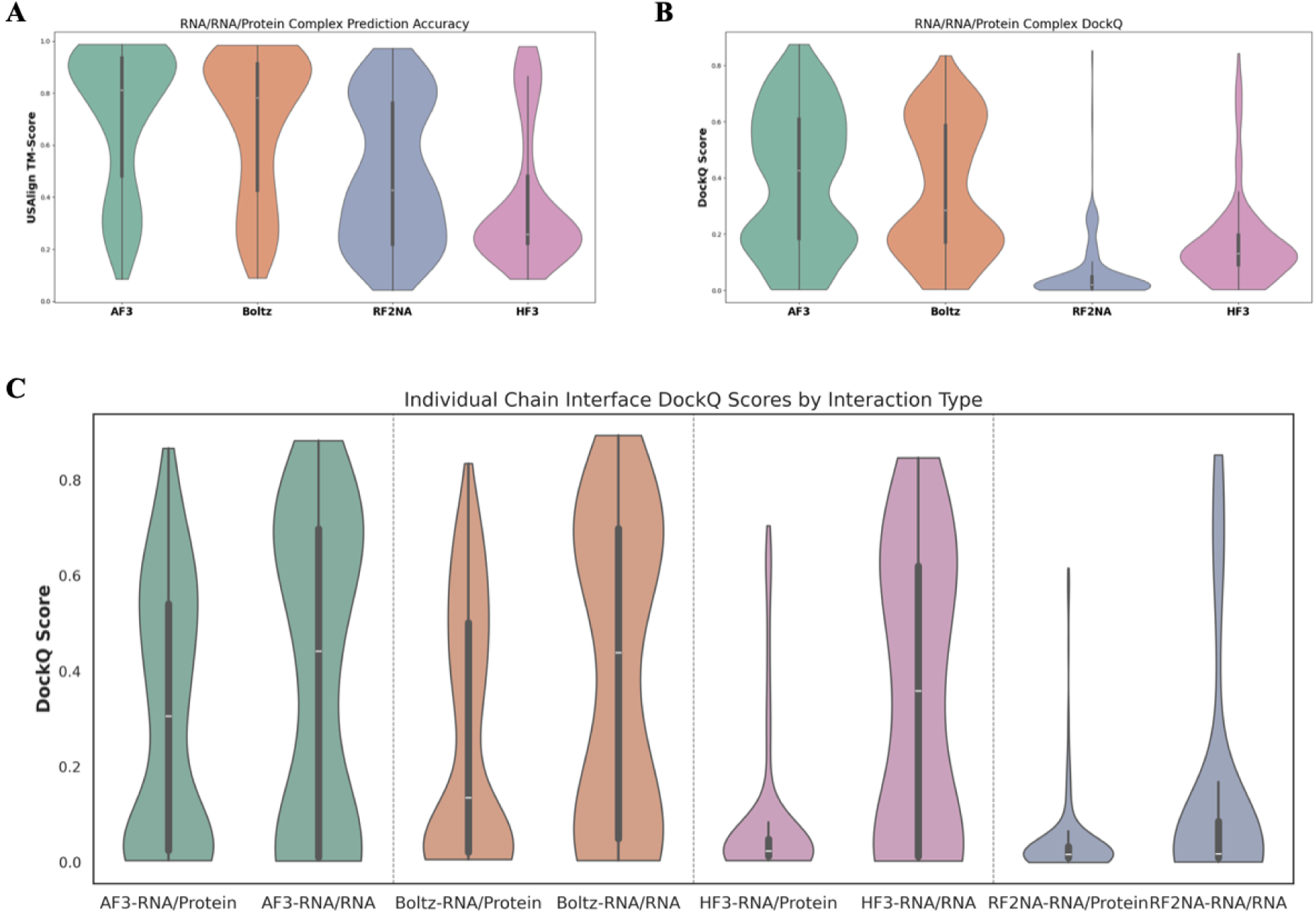
Performance comparison for different RNA prediction methods on RNA complexes. (A) The performance of RNA complex prediction models was evaluated using US-align. Each method corresponds to a separate violin plot. The horizontal line inside each violin plot represents the median, while the thin bar in the centre represents the interquartile range. Violin plots are shown in a descending order of TM-score in both cases. (B) Performance of RNA complex prediction models evaluated using DockQ score, following the same layout as in TM-score analysis. (C) DockQ score for individual pairs of chains (Interface DockQ, IF-DockQ). DockQ scores were calculated separately for RNA-protein (left) and RNA-RNA (right) complexes and shown as violin plots arranged in descending order based on the RNA-protein IF-DockQ scores.

To gain more insights, we analyzed individual RNA-RNA and RNA-protein interfaces within each complex by calculating DockQ scores for each pair of chains, i.e., the interface DockQ (IF-DockQ). The individual DockQ scores for RNA-RNA interfaces, compared with those for RNA-protein interfaces, suggest no significant difference in prediction accuracy between the two (Figure 6C; Table S9) for the best methods, while RNA-protein are worse than RNA-RNA predictions for both RFNA and HF3. AlphaFold3 shows the highest mean IF-DockQ score for RNA-protein (0.305), while Boltz-1 shows th highest mean IF-DockQ for RNA-RNA interfaces (0.393).

The relationships of the model qualities between different methods are shown in Figures 7A, 7B, and S9. AlphaFold3 outperformed Boltz-1 in some cases. A statistically significant difference was found for TM-Score (P = 3.85×10^-7^; Figure 7A), but no statistically significant difference was found for DockQ score (P = 1.05×10^-1^; Figure 7B).

**Figure 7.**
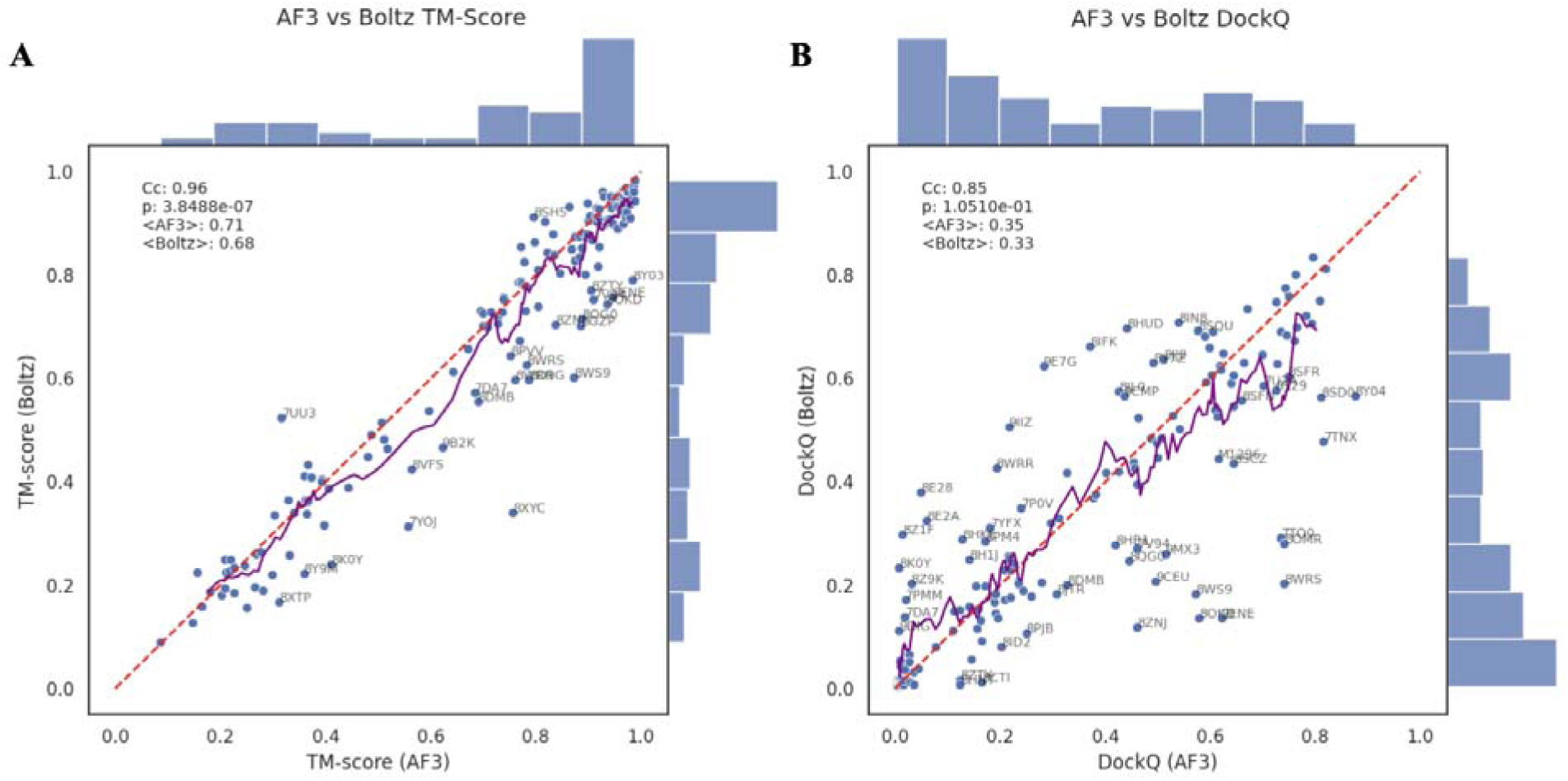
(A) Scatter plot showing correlation between AlphaFold3 and Boltz-1 TM-score for RNA complexes (C = 0.96, P = 3.85*10^-7^). A rolling average of the scores from the two methods is shown as a purple line. Bar plots o each side show the distribution of scores for each model separately (AlphaFold3 on the x-axis, Boltz on the y-axis). E) Scatter plot showing correlation between AlphaFold3 and Boltz-1 DockQ score for complexes (Cc = 0.85, P = 1.05*10^-1^).

Interface DockQ scores across all methods revealed that the same models consistently achieved the highest scores. The highest IF-DockQ between an RNA and a protein chain was achieved by HF3 in 8T29 (0.670), followed closely by AF3 in 8T29 (0.647). The highest IF-DockQ between two RNA chains was achieved by Boltz-1 in 7TNY (0.894), followed by AF3 in 7TNX (0.883). Notably, these highly accurate models are either tRNA-like RNAs or double-helical RNAs in protein complexes, likely benefiting from the relative simplicity of base-pair complementarity, which makes them easier to predict (Figure S10).

### Models with low DockQ scores and high TM-scores

To investigate whether accurate prediction of the overall complex fold also implies correct modeling of RNA-protein or RNA-RNA interfaces, we examined the relationship between interface quality and global structural accuracy. Specifically, for each method, we generated scatter plots comparing DockQ score against TM-score (Figure 8A-D). Across many complexes, we observe substantial differences between DockQ and TM-score, revealing that these two measures capture distinct aspects of prediction quality. In addition, this also indicates that for many models, although the overall fold of the complex is often modelled correctly, the binding interface is not predicted accurately. In such cases, TM-score exceeds 0.45, while the corresponding DockQ falls below the 0.23 threshold for acceptable interface accuracy. AF3 and Boltz-1 still show much higher agreement between these two metrics than HF3 and RF2NA. Consequently, the correlation between IF-DockQ and TM-score is weak (Figure 8 A-D).

**Figure 8.**
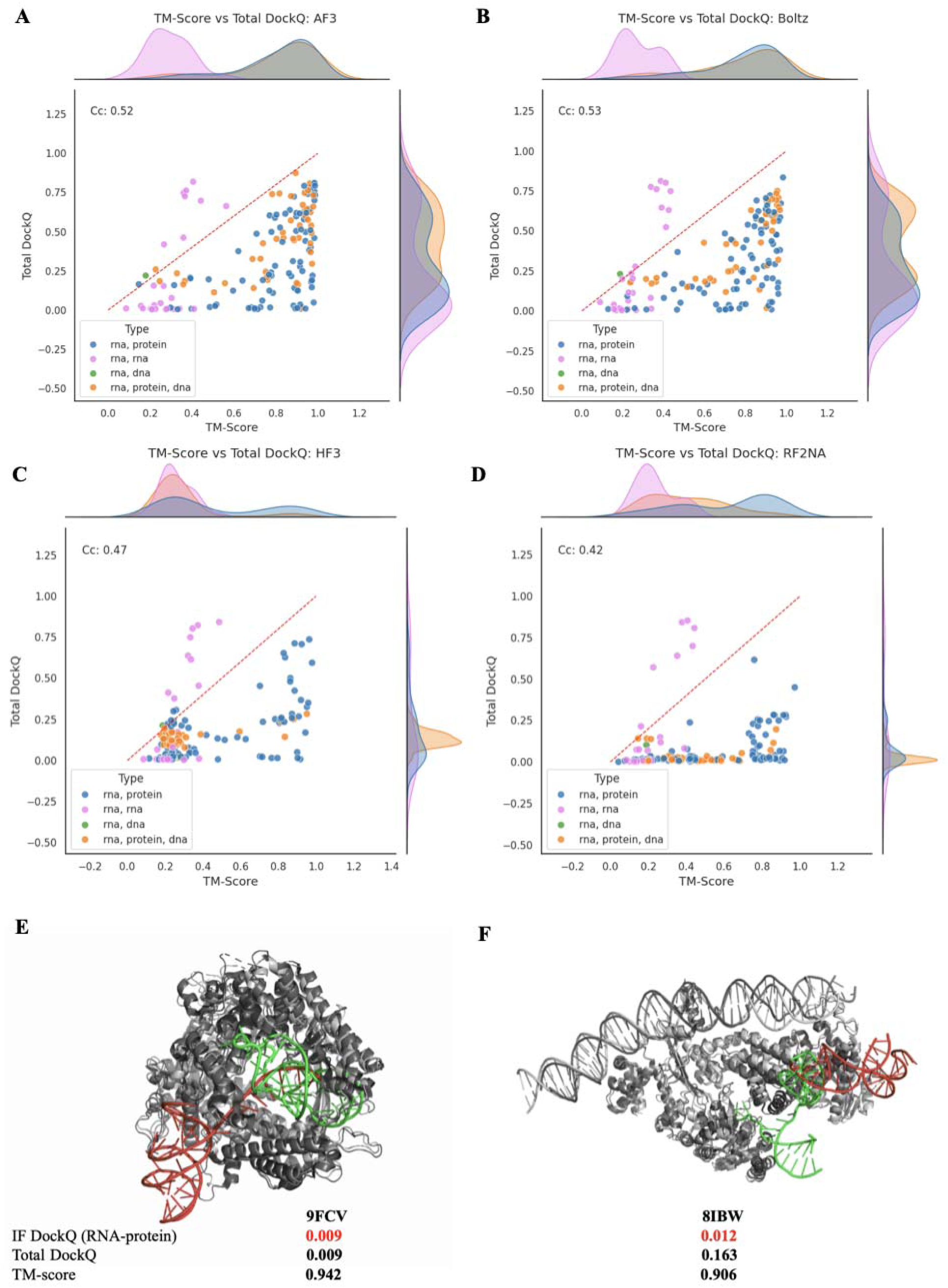
Scatter plots showing the correlation between TM score and DockQ. Points are colored based on the complex type (RNA-protein in blue, RNA-RNA in pink, RNA-DNA in green and RNA-protein-DNA in orange). Plots are presented for each RNA complex prediction method: (A) AlphaFold3, (B) Boltz-1, (C) HelixFold3, (D) RF2NA. (E) Structure of 9FCV predicted with AF3. (F) Structure of 8IBW predicted with AF3. In both examples, the reference RNA chain is shown in green, while the predicted misplaced RNA chain is colored red. TM-score is high, indicating an accurately modelled topology of complexes, while the IF-DockQ indicates incorrectly predicted binding sites.

The most pronounced example of this discrepancy is observed for the complex 9FCV, where both the protein and RNA components adopt accurate individual topologies, but the RNA is incorrectly positioned relative to the protein and binds at an incorrect binding site. This resulted in a very low DockQ score and an IF-DockQ between RNA and protein of only 0.009 (Figure 8E). The second-largest discrepancy was found in a model 8IN8, in which RNA chains were predicted to bind in the opposite orientation, so that both RNA chains that make up a double helix are bound to the wrong protein partners, resulting in a DockQ score of 0.163 and an IF-DockQ of 0.012 (Figure 8F).

Additionally, there are a few RNA-RNA complexes that show high DockQ scores, but very low TM-score. These structures are short RNA-RNA complexes (< 20 nt) where TM-score underevaluates structural accuracy, similarly as for single-chain RNAs. In total, there were 8 such structures: 7ECJ, 7ECK, 8HB1, 8YDC, 9BXE, 9CPD, 9CPJ and 9I8B.

### The reliability of pTM/ipTM confidence scores

Several methods, including AlphaFold3, Boltz-1, HelixFold3 and Chai-1, provide confidence metric alongside predicted structures in the form of predicted Template Modelling (pTM) scores for both monomeric and complex structures, and the interface predicted Template Modelling (ipTM) scores for macromolecular complexes. To assess the predictive reliability of these metrics, we compared them with the calculated TM-score for single-chain RNAs and RNA complexes, and with the DockQ score for RNA complexes (Figure 9 and S10).

**Figure 9.**
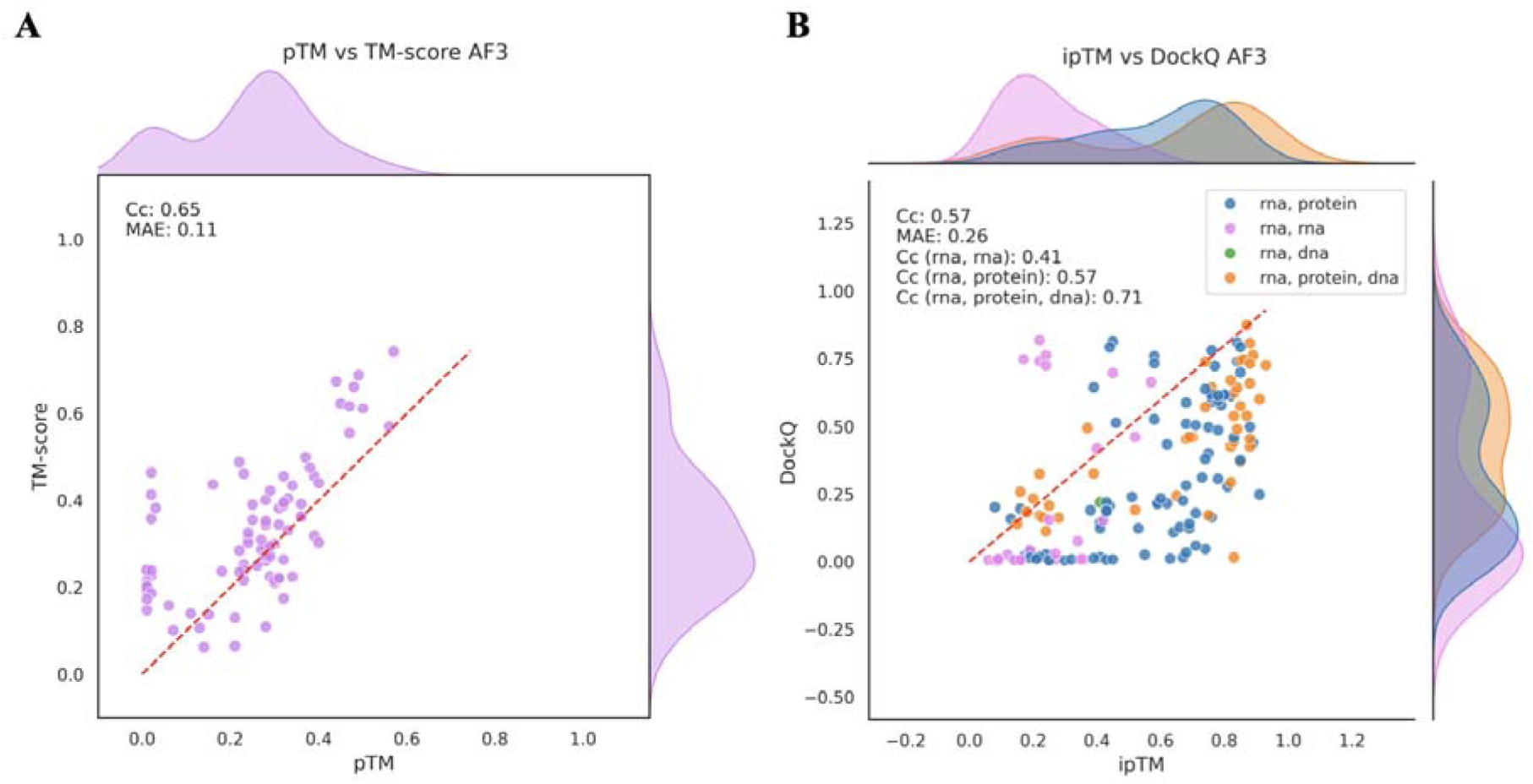
Evaluation of pTM/ipTM confidence scores. (A) Scatter plot between TM-score and pTM score for AlphaFold 3 and single-chain RNA models. (B) Scatter plot between DockQ score and ipTM score for AF3 and RNA complexes. RNA-RNA complexes are colored in pink, RNA-protein in blue and RNA-DNA in green and RNA-protein-DNA in orange.

For single-chain RNAs, the predicted TM scores (pTM) are generally low (pTM < 0.5) across all methods, except for Chai-1, which overestimated pTM scores (Figure S8E). The relationship between pTM and TM-score for AF3 is shown in Figure 8A, while the same plots for the other methods are provided in Figure S11. The overall correlation between pTM and TM-score remains strong, with a correlation coefficient of 0.65 and an MAE of 0.11 for AF3 (Figure 9A).

In contrast to single-chain RNAs, predicted interface TM scores (ipTM) for complexes, especially RNA-protein interactions, often exceed 0.6, indicating correct predictions, whereas DockQ values show that the models are incorrect (Figure 8B for AF3; Figure S10 for other methods). However, ipTM correlates moderately well with interface accuracy measured by DockQ, with the strongest correlation observed for Boltz-1 (Cc = 0.64), followed by HF3 (Cc = 0.61) and AF3 (Cc = 0.57). Notably, RNA-RNA complexes show consistently lower ipTM scores than RNA-protein complexes, suggesting that interface quality in RNA-RNA complexes is harder to estimate reliably. In the case of AF3, the mean ipTM for RNA-RNA complexes is 0.246, while RNA-protein complexes achieve a mean ipTM of 0.585 (Table S10). The difference of the DockQ scores between the categories is smaller (0.384 vs 0.305), as shown previously (Figure 6C, Table S9). Furthermore, mean chain ipTM for RNA chains (all interfaces where RNA chain interacts with any other chain) is 0.412, while mean chain ipTM for protein chains (all interfaces where protein chain interacts with any other chain) is higher (0.556), see Table S11. Therefore, the higher confidence observed for RNA-protein complexes arises from the presence of protein chains, whose structures are better learned by current methods, which increases overall confidence.

### Prediction accuracy dependence on AlphaFold3 training set

The presence of commonly found RNA motifs in the PDB raises the possibility that the benchmarked methods may perform well by recognizing frequently occurring patterns, rather than generalizing to novel or less common RNA structures. In other words, methods may primarily detect structural similarity to the training set.

As previously mentioned, our results show that double helical and cloverleaf structures are usually predicted with higher accuracy. However, these motifs are also among the most common in PDB. To assess the extent to which AlphaFold3’s prediction accuracy depends on prior exposure to similar RNA structures, we evaluated structural similarity between single-chain RNA benchmark targets and all RNA-containing entries in the AlphaFold3 training set.

For each RNA chain we computed TM-scores against all RNA chains in the training set. We retained the maximum TM-score per structure, corresponding to the highest structural similarity defined as maximum similarity to the training set (Figure 10A). We also defined four similarity categories using the following TM-score thresholds: <0.5, 0.25-0.5, 0.5–0.75, and 0.75 (Figure 10B). We observed that prediction accuracy, as measured by TM-score, generally increased with higher structural similarity to the training set. The mean TM-score is low for the category with the least similarity and increases gradually to over 0.5 for the category with the highest similarity to the training set. This suggests that AF3 tends to perform better when the target structure is more similar to motifs it encountered during training, highlighting the limitations of current methods in predicting unseen, structurally divergent RNAs. The same is observed for other methods and it is shown in Fig S12-13.

**Figure 10.**
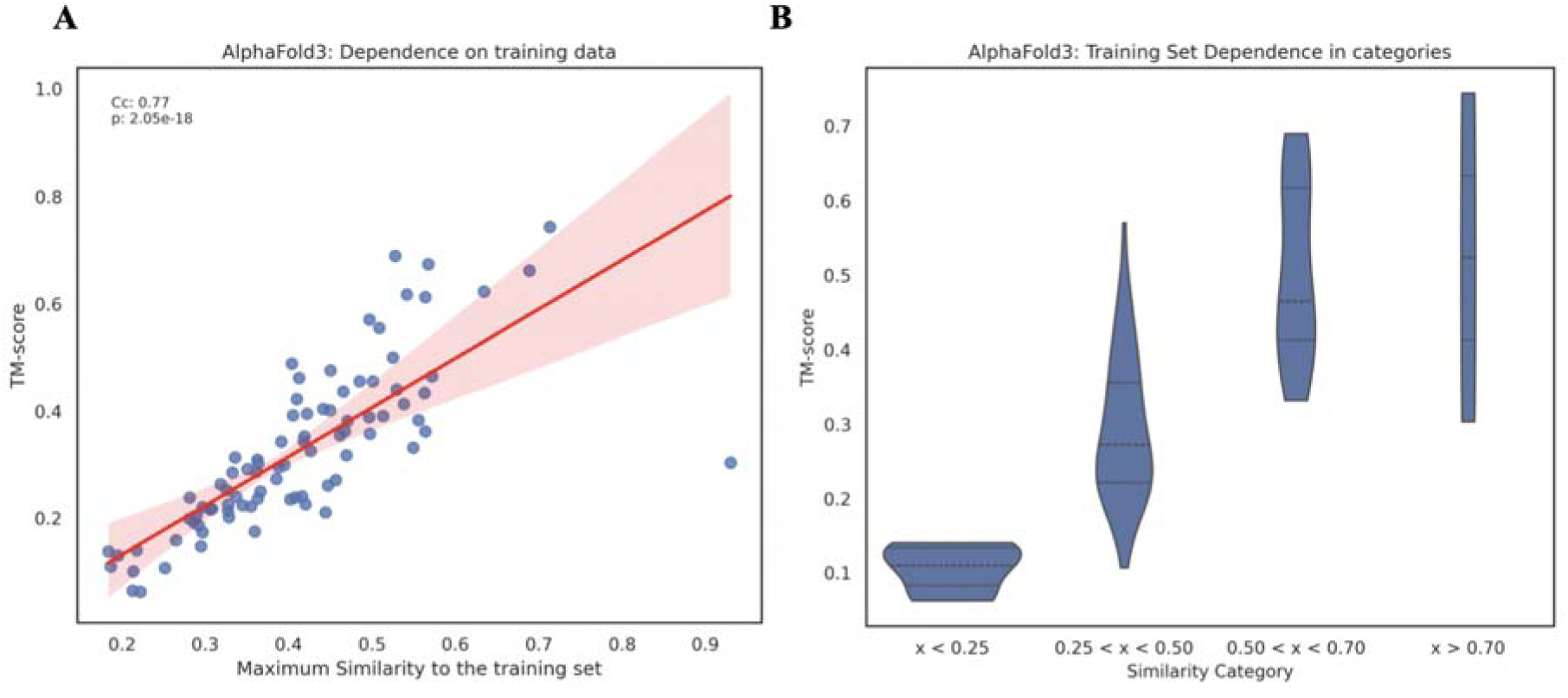
Similarity to the training set. (A) The relationship between prediction accuracy (TM-score) and maximum structural similarity to the AF3 training set for each single-chain RNA. (B) Prediction accuracy stratified by four training set similarity categories (TM-score: < 0.25, 0.2 5 - 0.5, 0.5 - 0.75, > 0.70).

## Conclusion

Advancements in deep learning have significantly impacted RNA structural biology, enabling the prediction of both monomeric RNA structures and RNA complexes. In this benchmark study, we systematically evaluated several state-of-the-art deep learning methods for RNA structure prediction, revealing their strengths and limitations. Our results suggest that while current methods show promising performance, particularly for well-characterized RNA motifs, substantial challenges remain.

Across all models, fewer than one-third achieve high accuracy according to our multi-metric success criteria. These models accurately predict predominantly RNAs structurally similar to previously solved RNA structures, such as L-shaped RNAs, RNAs dominated by simple loop motifs, and molecules with a high fraction of paired bases, including double-helical RNAs. In contrast, longer RNAs and those with more complex folds are consistently more difficult to predict accurately. This performance gap highlights a central limitation of current RNA structure prediction methods: their reliance on training data that does not cover the full diversity of RNA structural space, possibly due to the lack of diverse solved RNA structures deposited in the PDB and the dynamic nature of RNA. This is why the performance of those methods becomes accurate only on RNAs that resemble known folds, but struggle to generalize to novel structures. Similarly, a strong dependence of predictor performance on how closely an available template matched the target was also discussed in the CASP16 evaluation (10). Therefore, these findings underscore the need for broader, more structurally diverse RNA training datasets, as well as improved strategies for model selection and confidence estimation.

One of the most pronounced challenges identified in this benchmark is the accurate modelling of RNA-protein interactions. In many cases, both the RNA and protein chains individually adopt the correct global fold, yet the RNA is incorrectly bound to the wrong region of the protein surface. A likely explanation for this behaviour is bias in the training data, where the method may preferentially identify protein binding sites that are frequently involved in interactions in the PDB but are associated with different ligand or protein-protein interfaces rather than RNA.

Furthermore, while internal confidence metrics such as pTM and ipTM generally correlate with global and interface-level evaluation metrics, they should not be treated as definitive indicators of model quality, particularly for RNAs that lack close structural homologs. In several cases, accurate global folds receive low confidence scores, whereas high ipTM values in complexes, especially RNA-protein assemblies, may partly reflect the presence of well-learned protein components rather than true RNA interface accuracy. In the absence of an experimentally resolved structure or a closely related RNA family member with a known fold, we still recommend interpreting predicted RNA structures cautiously and, where possible, complementing them with experimental validation.

RNA is increasingly recognized as a versatile macromolecule with essential catalytic and regulatory functions, many of which are still unknown. We anticipate that growing interest in RNA biology will stimulate further experimental efforts towards solving complex RNA structures. Combined with innovations in techniques such as cryo-EM (2) and continued advances in machine learning (28), we are optimistic that RNA 3D structure prediction will evolve beyond simple motifs and toward accurately modelling the full complexity of RNA structural space.

## Acknowledgements

We thank the members of the Elofsson group for valuable input.

## Supplementary data

Supplementary data are available at NAR online.

## Conflict of interest

The authors declare no conflict of interest.

## Author Contributions

ML did all the computational work, wrote the final draft and the final version, and performed the analysis. AE obtained funding and helped with analysis and writing.

## Funding

AE was funded by the Vetenskapsrådet Grant No. 2021-03979 and the Knut and Alice Wallenberg Foundation, Grant No. 2022.0032. The computations and data handling were enabled by the supercomputing resource Berzelius, provided by the National Supercomputer Centre at Linköping University, the Knut and Alice Wallenberg Foundation, and SNIC, with grant Numbers. SNIC 2021/5-297 and Berzelius-2021-29.

## Data availability

Code used for running the methods and analyzing the main results is available from https://doi.org/10.5281/zenodo.18847731 and https://github.com/iammarcol/RNA-Benchmark

## Notes

https://github.com/iammarcol/RNA-Benchmark

### Competing Interest Statement

The authors have declared no competing interest.

### Summary of Updates

Update version with many changed/improvements. In short, we do find that the best methods accurately predict the structure of one third to slightly more than half of the structures/complexes. The performance is similar between the best methods (AlphaFold3 and Boltz-1) and only slighlty worse for other methods. However, we do note that basically all structures that can be predicted accurately share structural similarity (not necessarily homology though) with entries in PDB. Thie means that curren tstate of the art methods can not be used to predict the structure of "novel" RNA-molecules. Further, the ability to estimate the accuracy of the predictions (the pTM and ipTM values) is significantly worse than for proteins. This limits the ability to do large scale structure prediction studies and we do believe this is an important area for improvements

