## Supplementary Information for "Limits of deep-learning-based RNA prediction methods"

### Supplementary Tables

**Table S1.** TM-score for the whole benchmarked dataset of single-chain RNA molecules across all tested deep learning methods for single-chain RNA prediction.

| PDB | AF3 | Boltz | Chai | HF3 | NuFold | RF2NA | RhoFold | trRosettaRNA |
|---|---|---|---|---|---|---|---|---|
| **7EXY** | 0.22726 | 0.27096 | 0.20765 | 0.25169 | 0.26408 | 0.21924 | 0.16804 | 0.15878 |
| **7KUB** | 0.36302 | 0.39686 | 0.33353 | 0.35902 | 0.4023 | 0.4335 | 0.33309 | 0.38691 |
| **7KUC** | 0.46535 | 0.33929 | 0.33573 | 0.36668 | 0.41476 | 0.40323 | 0.26203 | 0.36668 |
| **7KUD** | 0.38398 | 0.27041 | 0.30498 | 0.35551 | 0.34221 | 0.20897 | 0.12448 | 0.38897 |
| **7MKT** | 0.06505 | 0.10619 | 0.11349 | 0.12008 | 0.08987 | 0.07464 | 0.06848 | 0.09654 |
| **7PS8** | 0.13037 | 0.10516 | 0.14895 | 0.13334 | 0.09658 | 0.13584 | 0.1169 | 0.12639 |
| **7Q48** | 0.10714 | 0.12714 | 0.12852 | 0.13898 | 0.09031 | 0.13405 | 0.11763 | 0.13234 |
| **7Q6L** | 0.10097 | 0.14818 | 0.12497 | 0.14837 | 0.11873 | 0.14177 | 0.12734 | 0.15612 |
| **7QA2** | 0.14075 | 0.15234 | 0.11873 | 0.12574 | 0.12327 | 0.11432 | 0.1641 | 0.10928 |
| **7RWR** | 0.38245 | 0.44787 | 0.27836 | 0.28942 | 0.37568 | 0.36085 | 0.39044 | 0.39621 |
| **7SHX** | 0.17544 | 0.24919 | 0.17696 | 0.26902 | 0.17875 | 0.19857 | 0.19728 | 0.19667 |
| **7SXP** | 0.06271 | 0.23686 | 0.13791 | 0.0447 | 0.12365 | 0.09124 | 0.06944 | 0.17309 |
| **7UGA** | 0.40508 | 0.32635 | 0.3167 | 0.31472 | 0.42513 | 0.33186 | 0.33502 | 0.33883 |
| **7UMC** | 0.34501 | 0.41391 | 0.31398 | 0.29232 | 0.31915 | 0.4994 | 0.36128 | 0.29601 |
| **7UMD** | 0.29226 | 0.29686 | 0.34879 | 0.21713 | 0.31574 | 0.2611 | 0.3279 | 0.16665 |
| **7UME** | 0.39126 | 0.36121 | 0.40673 | 0.22378 | 0.38316 | 0.30098 | 0.27976 | 0.14482 |
| **7V06** | 0.31867 | 0.389 | 0.48251 | 0.31508 | 0.35875 | 0.31852 | 0.3823 | 0.30012 |
| **7WIA** | 0.38924 | 0.3466 | 0.32442 | 0.26068 | 0.42405 | 0.39318 | 0.30375 | 0.38543 |
| **8BTZ** | 0.28753 | 0.22254 | 0.54226 | 0.25987 | 0.3149 | 0.26744 | 0.3763 | 0.21728 |
| **8BU8** | 0.2378 | 0.35885 | 0.23294 | 0.35399 | 0.27524 | 0.26255 | 0.15836 | 0.31081 |

| | | | | | | | | |
|---|---|---|---|---|---|---|---|---|
| **8BWT** | 0.22597 | 0.21981 | 0.24246 | 0.21498 | 0.20018 | 0.21143 | 0.22829 | 0.17153 |
| **8CLR** | 0.24165 | 0.22713 | 0.25075 | 0.27882 | 0.30522 | 0.28221 | 0.23224 | 0.20346 |
| **8CQ1** | 0.45643 | 0.42347 | 0.38626 | 0.38878 | 0.41691 | 0.43917 | 0.36146 | 0.40835 |
| **8FB3** | 0.36314 | 0.37594 | 0.25761 | 0.26562 | 0.58616 | 0.30276 | 0.46508 | 0.2659 |
| **8FCS** | 0.67471 | 615 | 0.59979 | 0.43196 | 0.63414 | 0.60651 | 0.62008 | 0.50856 |
| **8FZA** | 0.26184 | 0.19618 | 0.18728 | 0.21402 | 0.25951 | 0.24406 | 0.17132 | 0.27127 |
| **8HB8** | 0.22502 | 0.19319 | 0.29295 | 0.19791 | 0.27355 | 0.22435 | 0.21816 | 0.14942 |
| **8I43** | 0.19923 | 0.19486 | 0.16197 | 0.26151 | 0.20185 | 0.22929 | 0.15882 | 0.14755 |
| **8I44** | 0.21506 | 0.21954 | 0.17371 | 0.15447 | 0.20255 | 0.16523 | 0.15596 | 0.13993 |
| **8I45** | 0.20611 | 0.19284 | 0.16642 | 0.20113 | 0.18629 | 0.21105 | 0.15649 | 0.13365 |
| **8I46** | 0.19182 | 0.21902 | 0.18004 | 0.24284 | 0.2308 | 0.18263 | 0.19841 | 0.14397 |
| **8ITS** | 0.3098 | 0.26224 | 0.28633 | 0.29085 | 0.25766 | 0.30097 | 0.21876 | 0.17217 |
| **8JHP** | 0.21709 | 0.25444 | 0.24458 | 0.22424 | 0.24537 | 0.17925 | 0.16361 | 0.1402 |
| **8K2Z** | 0.55552 | 0.58768 | 0.34971 | 0.29386 | 0.44706 | 0.27612 | 0.35743 | 0.13557 |
| **8K30** | 0.61317 | 0.37553 | 0.53594 | 0.3084 | 0.63489 | 0.33291 | 0.40124 | 0.53919 |
| **8K8B** | 0.23959 | 0.20711 | 0.17385 | 0.20575 | 0.16605 | 0.18501 | 0.16353 | 0.18352 |
| **8KEB** | 0.66235 | 0.60309 | 0.31577 | 0.69134 | 0.71253 | 0.27968 | 0.59691 | 0.1606 |
| **8Q4O** | 0.13807 | 0.11463 | 0.10855 | 0.13689 | 0.14232 | 0.12586 | 0.10461 | 0.10659 |
| **8Q6N** | 0.33247 | 0.54981 | 0.40027 | 0.44513 | 0.54175 | 0.34414 | 0.3297 | 0.12914 |
| **8QO2** | 0.46272 | 0.24538 | 0.22837 | 0.20328 | 0.31543 | 0.27924 | 0.44752 | 0.28936 |
| **8QO3** | 0.48945 | 0.41034 | 0.39579 | 0.37517 | 0.5326 | 0.3651 | 0.43298 | 0.36604 |
| **8QO4** | 0.40196 | 0.41335 | 0.27552 | 0.33193 | 0.38195 | 0.36692 | 0.43898 | 0.29529 |
| **8QO5** | 0.42299 | 0.31311 | 0.23883 | 0.36726 | 0.29979 | 0.48055 | 0.26341 | 0.21046 |
| **8SCF** | 0.45615 | 0.41437 | 0.35514 | 0.39699 | 0.41473 | 0.45921 | 0.29352 | 0.11683 |
| **8SCH** | 0.34403 | 0.29319 | 0.3242 | 0.29693 | 0.23178 | 0.32097 | 0.3039 | 0.2726 |
| **8T5O** | 0.25107 | 0.30955 | 0.2834 | 0.15921 | 0.21788 | 0.25267 | 0.3091 | 0.21043 |
| **8TNS** | 0.1101 | 0.14566 | 0.10459 | 0.10507 | 0.13141 | 0.07363 | 0.09314 | 0.08636 |
| **8TVZ** | 0.25297 | 0.44026 | 0.41025 | 0.33252 | 0.24662 | 0.17997 | 0.11467 | 0.21138 |
| **8UPT** | 0.62407 | 0.66113 | 0.40738 | 0.21266 | 0.6028 | 0.38416 | 0.46457 | 0.49638 |
| **8UTG** | 0.15968 | 0.16531 | 0.05151 | 0.12801 | 0.14738 | 0.13584 | 0.09691 | 0.13591 |
| **8UYG** | 0.39278 | 0.36566 | 0.30983 | 0.30681 | 0.29237 | 0.33127 | 0.34361 | 0.22807 |
| **8UYM** | 0.39602 | 0.40098 | 0.27704 | 0.38417 | 0.27121 | 0.39409 | 0.32686 | 0.31631 |

| | | | | | | | | |
|---|---|---|---|---|---|---|---|---|
| **8UYP** | 0.43726 | 0.31299 | 0.22662 | 0.38017 | 0.37182 | 0.42463 | 0.32979 | 0.20298 |
| **8V1H** | 0.57092 | 0.59434 | 0.56451 | 0.58333 | 0.55012 | 0.46147 | 0.49749 | 0.13601 |
| **8V1I** | 0.61837 | 0.56803 | 0.42743 | 0.29223 | 0.5797 | 0.37802 | 0.40787 | 0.48961 |
| **8VCI** | 0.27375 | 0.36171 | 0.23514 | 0.35249 | 0.28861 | 0.29306 | 0.27583 | 0.32766 |
| **8VT5** | 0.68961 | 0.67695 | 0.46968 | 0.40826 | 0.52719 | 0.5076 | 0.48161 | 0.54258 |
| **8XZL** | 0.27199 | 0.40012 | 0.30423 | 0.2601 | 0.27101 | 0.36679 | 0.25009 | 0.20187 |
| **8ZAU** | 0.35335 | 0.29659 | 0.3289 | 0.33727 | 0.3821 | 0.26212 | 0.5033 | 0.25314 |
| **9BLM** | 0.44052 | 0.51669 | 0.37959 | 0.23524 | 0.56686 | 0.44748 | 0.47366 | 0.28217 |
| **9BUN** | 0.21178 | 0.38283 | 0.37039 | 0.36321 | 0.32323 | 0.286 | 0.29476 | 0.28951 |
| **9DE6** | 0.43479 | 0.26276 | 0.55175 | 0.53952 | 0.53482 | 0.59007 | 0.35804 | 0.43245 |
| **9DE7** | 0.50035 | 0.44777 | 0.48962 | 0.50675 | 0.49865 | 0.30042 | 0.39677 | 0.47788 |
| **9ECQ** | 0.35831 | 0.50386 | 0.37724 | 0.27222 | 0.42793 | 0.35474 | 0.24676 | 0.35127 |
| **9ECR** | 0.41407 | 0.37967 | 0.4179 | 0.29519 | 0.3495 | 0.44445 | 0.27964 | 0.31863 |
| **9EOW** | 0.21778 | 0.20014 | 0.16285 | 0.18449 | 0.1944 | 0.21459 | 0.17013 | 0.13366 |
| **9EWW** | 0.2224 | 0.26513 | 0.1944 | 0.17392 | 0.18445 | 0.1918 | 0.14388 | 0.18433 |
| **9FM4** | 0.24104 | 0.2416 | 0.20345 | 0.2167 | 0.21894 | 0.20931 | 0.16578 | 0.23808 |
| **9FO8** | 0.22161 | 0.23082 | 0.1992 | 0.17442 | 0.23475 | 0.19325 | 0.17154 | 0.19494 |
| **9FO9** | 0.26401 | 0.24704 | 0.2359 | 0.25152 | 0.25481 | 0.20034 | 0.21836 | 0.21747 |
| **9G7C** | 0.31392 | 0.22116 | 0.22217 | 0.21041 | 0.23659 | 0.20471 | 0.25365 | 0.20089 |
| **9IO0** | 0.20246 | 0.24801 | 0.18491 | 0.19481 | 0.19119 | 0.17503 | 0.16545 | 0.22239 |
| **9IO1** | 0.14843 | 0.21158 | 0.14867 | 0.20285 | 0.15892 | 0.19405 | 0.16813 | 0.20374 |
| **9IOR** | 0.23935 | 0.2393 | 0.24999 | 0.22025 | 0.23692 | 0.21245 | 0.16596 | 0.14246 |
| **9IOS** | 0.18702 | 0.22048 | 0.19625 | 0.1721 | 0.18025 | 0.19107 | 0.12374 | 0.11185 |
| **9IOU** | 0.17398 | 0.17086 | 0.20248 | 0.21529 | 0.18137 | 0.19258 | 0.15022 | 0.15908 |
| **R1205** | 0.30023 | 0.25182 | 0.269 | 0.34961 | 0.32268 | 0.34767 | 0.24638 | 0.25095 |
| **R1209** | 0.3027 | 0.30327 | 0.25469 | 0.23275 | 0.27681 | 0.3004 | 0.23844 | 0.25086 |
| **R1211** | 0.47669 | 0.41407 | 0.41496 | 0.50952 | 0.45052 | 0.34751 | 0.30872 | 0.33897 |
| **R1212** | 0.23588 | 0.239 | 0.24179 | 0.25041 | 0.23643 | 0.2133 | 0.26621 | 0.23704 |
| **R1248** | 0.28572 | 0.22022 | 0.23926 | 0.28349 | 0.25438 | 0.20922 | 0.27824 | 0.2729 |
| **R1263** | 0.304 | 0.49274 | 0.27553 | 0.48693 | 0.66333 | 0.92072 | 0.48397 | 0.7898 |
| **R1271** | 0.74432 | 0.72539 | 0.62939 | 0.66229 | 0.66617 | 0.61516 | 0.57549 | 0.1657 |
| **R1288** | 0.29578 | 0.30832 | 0.24632 | 0.2532 | 0.34882 | 0.29095 | 0.25044 | 0.18367 |

| | | | | | | | | |
|---|---|---|---|---|---|---|---|---|
| **R1293** | 0.35571 | 0.36727 | 0.45625 | 0.40129 | 0.31029 | 0.24329 | 0.23932 | 0.38649 |
| **R1296** | 0.32625 | 0.29787 | 0.26851 | 0.33956 | 0.3388 | 0.35016 | 0.29936 | 0.2846 |

**Table S2.** GDT-TS for the whole benchmarked dataset of single-chain RNA molecules across all tested deep learning methods for single-chain RNA prediction.

| PDB | AF3 | Boltz | Chai | HF3 | NuFold | RF2NA | RhoFold | trRosettaRNA |
|---|---|---|---|---|---|---|---|---|
| **7EXY** | 0.64 | 0.75 | 0.69 | 0.62 | 0.66 | 0.69 | 0.7 | 0.45 |
| **7KUB** | 0.22 | 0.24 | 0.23 | 0.23 | 0.25 | 0.25 | 0.23 | 0.2 |
| **7KUC** | 0.88 | 0.86 | 0.88 | 0.84 | 0.94 | 0.91 | 0.88 | 0.92 |
| **7KUD** | 0.86 | 0.83 | 0.81 | 0.94 | 0.9 | 0.81 | 0.79 | 0.58 |
| **7MKT** | 0.3 | 0.29 | 0.26 | 0.33 | 0.26 | 0.22 | 0.25 | 0.25 |
| **7PS8** | 0.32 | 0.45 | 0.28 | 0.32 | 0.29 | 0.3 | 0.29 | 0.3 |
| **7Q48** | 0.31 | 0.38 | 0.32 | 0.3 | 0.31 | 0.36 | 0.32 | 0.34 |
| **7Q6L** | 0.32 | 0.39 | 0.32 | 0.35 | 0.31 | 0.33 | 0.32 | 0.32 |
| **7QA2** | 0.34 | 0.43 | 0.33 | 0.34 | 0.31 | 0.33 | 0.33 | 0.34 |
| **7RWR** | 0.64 | 0.7 | 0.57 | 0.48 | 0.64 | 0.6 | 0.62 | 0.66 |
| **7SHX** | 0.2 | 0.21 | 0.2 | 0.2 | 0.19 | 0.19 | 0.19 | 0.16 |
| **7SXP** | 0.37 | 0.54 | 0.38 | 0.47 | 0.34 | 0.34 | 0.36 | 0.32 |
| **7UGA** | 0.6 | 0.47 | 0.51 | 0.55 | 0.56 | 0.59 | 0.53 | 0.52 |
| **7UMC** | 0.4 | 0.51 | 0.36 | 0.34 | 0.39 | 0.43 | 0.45 | 0.34 |
| **7UMD** | 0.48 | 0.57 | 0.59 | 0.38 | 0.49 | 0.51 | 0.55 | 0.42 |
| **7UME** | 0.82 | 0.78 | 0.8 | 0.64 | 0.79 | 0.8 | 0.74 | 0.54 |
| **7V06** | 0.57 | 0.61 | 0.78 | 0.64 | 0.59 | 0.62 | 0.64 | 0.46 |
| **7WIA** | 0.24 | 0.24 | 0.25 | 0.19 | 0.26 | 0.25 | 0.24 | 0.22 |
| **8BTZ** | 0.17 | 0.13 | 0.34 | 0.19 | 0.21 | 0.16 | 0.22 | 0.12 |
| **8BU8** | 0.14 | 0.2 | 0.12 | 0.16 | 0.13 | 0.13 | 0.05 | 0.14 |
| **8BWT** | 0.66 | 0.65 | 0.69 | 0.67 | 0.6 | 0.62 | 0.66 | 0.53 |
| **8CLR** | 0.91 | 0.88 | 0.88 | 0.86 | 0.91 | 0.86 | 0.88 | 0.77 |
| **8CQ1** | 0.7 | 0.71 | 0.68 | 0.69 | 0.69 | 0.7 | 0.64 | 0.67 |
| **8FB3** | 0.71 | 0.76 | 0.26 | 0.57 | 0.92 | 0.53 | 0.86 | 0.68 |
| **8FCS** | 0.74 | 0.69 | 0.66 | 0.49 | 0.72 | 0.69 | 0.69 | 0.62 |
| **8FZA** | 0.75 | 0.46 | 0.47 | 0.73 | 0.73 | 0.45 | 0.65 | 0.76 |
| **8HB8** | 0.35 | 0.24 | 0.4 | 0.28 | 0.34 | 0.28 | 0.33 | 0.2 |
| **8I43** | 0.58 | 0.6 | 0.6 | 0.67 | 0.59 | 0.62 | 0.58 | 0.58 |

| | | | | | | | | |
|---|---|---|---|---|---|---|---|---|
| **8I44** | 0.67 | 0.67 | 0.63 | 0.7 | 0.66 | 0.64 | 0.62 | 0.38 |
| **8I45** | 0.76 | 0.71 | 0.7 | 0.71 | 0.7 | 0.72 | 0.67 | 0.42 |
| **8I46** | 0.79 | 0.71 | 0.68 | 0.75 | 0.76 | 0.7 | 0.62 | 0.72 |
| **8ITS** | 0.4 | 0.35 | 0.41 | 0.38 | 0.39 | 0.4 | 0.4 | 0.22 |
| **8JHP** | 0.56 | 0.61 | 0.69 | 0.49 | 0.57 | 0.59 | 0.56 | 0.58 |
| **8K2Z** | 0.71 | 0.75 | 0.51 | 0.43 | 0.64 | 0.42 | 0.56 | 0.23 |
| **8K30** | 0.8 | 0.51 | 0.69 | 0.38 | 0.79 | 0.45 | 0.65 | 0.72 |
| **8K8B** | 0.63 | 0.67 | 0.57 | 0.62 | 0.6 | 0.62 | 0.58 | 0.4 |
| **8KEB** | 0.76 | 0.69 | 0.37 | 0.77 | 0.79 | 0.33 | 0.67 | 0.22 |
| **8Q4O** | 0.34 | 0.32 | 0.3 | 0.32 | 0.27 | 0.3 | 0.3 | 0.33 |
| **8Q6N** | 0.57 | 0.78 | 0.59 | 0.66 | 0.8 | 0.59 | 0.5 | 0.27 |
| **8QO2** | 0.07 | 0.07 | 0.06 | 0.06 | 0.07 | 0.07 | 0.07 | 0.07 |
| **8QO3** | 0.37 | 0.35 | 0.35 | 0.34 | 0.39 | 0.28 | 0.38 | 0.28 |
| **8QO4** | 0.31 | 0.26 | 0.2 | 0.22 | 0.28 | 0.28 | 0.31 | 0.2 |
| **8QO5** | 0.33 | 0.21 | 0.19 | 0.26 | 0.19 | 0.32 | 0.21 | 0.17 |
| **8SCF** | 0.82 | 0.82 | 0.78 | 0.82 | 0.78 | 0.81 | 0.75 | 0.3 |
| **8SCH** | 0.42 | 0.39 | 0.38 | 0.28 | 0.35 | 0.41 | 0.38 | 0.35 |
| **8T5O** | 0.19 | 0.26 | 0.19 | 0.15 | 0.21 | 0.19 | 0.2 | 0.16 |
| **8TNS** | 0.31 | 0.29 | 0.34 | 0.28 | 0.3 | 0.21 | 0.24 | 0.26 |
| **8TVZ** | 0.11 | 0.17 | 0.15 | 0.12 | 0.1 | 0.1 | 0.02 | 0.1 |
| **8UPT** | 0.16 | 0.16 | 0.16 | 0.15 | 0.14 | 0.14 | 0.13 | 0.14 |
| **8UTG** | 0.33 | 0.48 | 0.34 | 0.39 | 0.34 | 0.32 | 0.33 | 0.3 |
| **8UYG** | 0.29 | 0.31 | 0.13 | 0.23 | 0.25 | 0.28 | 0.25 | 0.2 |
| **8UYM** | 0.36 | 0.33 | 0.24 | 0.33 | 0.23 | 0.28 | 0.26 | 0.25 |
| **8UYP** | 0.35 | 0.25 | 0.2 | 0.31 | 0.28 | 0.33 | 0.26 | 0.15 |
| **8V1H** | 0.19 | 0.18 | 0.19 | 0.18 | 0.2 | 0.18 | 0.18 | 0.12 |
| **8V1I** | 0.78 | 0.75 | 0.55 | 0.35 | 0.73 | 0.51 | 0.59 | 0.64 |
| **8VCI** | 0.23 | 0.28 | 0.22 | 0.29 | 0.22 | 0.21 | 0.25 | 0.28 |
| **8VT5** | 0.8 | 0.8 | 0.59 | 0.52 | 0.72 | 0.63 | 0.64 | 0.67 |
| **8XZL** | 0.39 | 0.59 | 0.42 | 0.38 | 0.43 | 0.49 | 0.43 | 0.32 |
| **8ZAU** | 0.39 | 0.34 | 0.34 | 0.36 | 0.38 | 0.3 | 0.6 | 0.23 |
| **9BLM** | 0.55 | 0.62 | 0.33 | 0.31 | 0.7 | 0.52 | 0.55 | 0.31 |

| | | | | | | | | |
|---|---|---|---|---|---|---|---|---|
| **9BUN** | 0.36 | 0.55 | 0.54 | 0.5 | 0.46 | 0.45 | 0.43 | 0.45 |
| **9DE6** | 0.33 | 0.34 | 0.45 | 0.47 | 0.43 | 0.52 | 0.36 | 0.4 |
| **9DE7** | 0.46 | 0.46 | 0.43 | 0.45 | 0.45 | 0.31 | 0.38 | 0.43 |
| **9ECQ** | 0.87 | 0.9 | 0.85 | 0.78 | 0.85 | 0.82 | 0.76 | 0.82 |
| **9ECR** | 0.64 | 0.67 | 0.67 | 0.59 | 0.64 | 0.66 | 0.69 | 0.64 |
| **9EOW** | 0.33 | 0.32 | 0.3 | 0.32 | 0.32 | 0.33 | 0.3 | 0.26 |
| **9EWW** | 0.2 | 0.2 | 0.09 | 0.16 | 0.17 | 0.18 | 0.13 | 0.16 |
| **9FM4** | 0.69 | 0.68 | 0.63 | 0.66 | 0.66 | 0.64 | 0.62 | 0.67 |
| **9FO8** | 0.6 | 0.62 | 0.53 | 0.58 | 0.62 | 0.61 | 0.59 | 0.62 |
| **9FO9** | 0.54 | 0.55 | 0.52 | 0.58 | 0.6 | 0.55 | 0.57 | 0.64 |
| **9G7C** | 0.18 | 0.15 | 0.13 | 0.14 | 0.16 | 0.13 | 0.16 | 0.15 |
| **9IO0** | 0.54 | 0.54 | 0.55 | 0.53 | 0.54 | 0.53 | 0.58 | 0.54 |
| **9IO1** | 0.59 | 0.53 | 0.51 | 0.58 | 0.55 | 0.57 | 0.47 | 0.55 |
| **9IOR** | 0.59 | 0.53 | 0.57 | 0.57 | 0.57 | 0.59 | 0.51 | 0.54 |
| **9IOS** | 0.5 | 0.51 | 0.49 | 0.46 | 0.51 | 0.53 | 0.42 | 0.5 |
| **9IOU** | 0.49 | 0.51 | 0.47 | 0.49 | 0.51 | 0.5 | 0.41 | 0.49 |
| **R1205** | 0.4 | 0.38 | 0.32 | 0.43 | 0.36 | 0.4 | 0.31 | 0.31 |
| **R1209** | 0.33 | 0.33 | 0.29 | 0.31 | 0.3 | 0.32 | 0.3 | 0.3 |
| **R1211** | 0.48 | 0.44 | 0.41 | 0.48 | 0.44 | 0.36 | 0.29 | 0.33 |
| **R1212** | 0.23 | 0.21 | 0.25 | 0.23 | 0.24 | 0.18 | 0.21 | 0.24 |
| **R1248** | 0.06 | 0.05 | 0.06 | 0.05 | 0.06 | 0.05 | 0.06 | 0.06 |
| **R1263** | 0.41 | 0.62 | 0.38 | 0.56 | 0.78 | 0.98 | 0.58 | 0.86 |
| **R1271** | 0.76 | 0.76 | 0.69 | 0.7 | 0.7 | 0.7 | 0.62 | 0.22 |
| **R1288** | 0.48 | 0.49 | 0.4 | 0.28 | 0.47 | 0.5 | 0.49 | 0.21 |
| **R1293** | 0.33 | 0.44 | 0.42 | 0.37 | 0.31 | 0.23 | 0.24 | 0.42 |
| **R1296** | 0.38 | 0.35 | 0.31 | 0.39 | 0.37 | 0.39 | 0.3 | 0.31 |

**Table S3.** INF-ALL for the whole benchmarked dataset of single-chain RNA molecules across all tested deep learning methods for single-chain RNA prediction.

| PDB | AF3 | Boltz | Chai | HF3 | NuFold | RF2NA | RhoFold | trRosettaRNA |
|---|---|---|---|---|---|---|---|---|
| 7EXY | 0.84 | 0.81 | 0.88 | 0.87 | 0.76 | 0.73 | 0.84 | 0.27 |
| 7KUB | 0.88 | 0.81 | 0.86 | 0.85 | 0.78 | 0.69 | 0.64 | 0.63 |
| 7KUC | 0.97 | 0.95 | 0.86 | 0.95 | 0.92 | 0.97 | 0.82 | 0.92 |
| 7KUD | 0.82 | 0.76 | 0.9 | 0.8 | 0.74 | 0.77 | 0.85 | 0.77 |
| 7MKT | 0.0 | 0.0 | 0.0 | 0.0 | 0.0 | 0.0 | 0.0 | 0.0 |
| 7PS8 | 0.25 | 0.31 | 0.35 | 0.21 | 0.26 | 0.33 | 0.23 | 0.26 |
| 7Q48 | 0.33 | 0.45 | 0.49 | 0.34 | 0.33 | 0.38 | 0.37 | 0.36 |
| 7Q6L | 0.31 | 0.36 | 0.36 | 0.32 | 0.3 | 0.45 | 0.36 | 0.28 |
| 7QA2 | 0.4 | 0.49 | 0.46 | 0.46 | 0.27 | 0.42 | 0.45 | 0.36 |
| 7RWR | 0.67 | 0.68 | 0.69 | 0.62 | 0.74 | 0.69 | 0.69 | 0.75 |
| 7SHX | 0.59 | 0.7 | 0.62 | 0.65 | 0.68 | 0.64 | 0.62 | 0.47 |
| 7SXP | 0.3 | 0.43 | 0.33 | 0.24 | 0.25 | 0.34 | 0.44 | 0.36 |
| 7UGA | 0.9 | 0.85 | 0.84 | 0.89 | 0.83 | 0.77 | 0.84 | 0.81 |
| 7UMC | 0.82 | 0.83 | 0.7 | 0.83 | 0.73 | 0.75 | 0.78 | 0.64 |
| 7UMD | 0.81 | 0.82 | 0.81 | 0.71 | 0.68 | 0.78 | 0.75 | 0.55 |
| 7UME | 0.82 | 0.81 | 0.77 | 0.8 | 0.83 | 0.76 | 0.78 | 0.47 |
| 7V06 | 0.72 | 0.74 | 0.83 | 0.8 | 0.62 | 0.73 | 0.72 | 0.49 |
| 7WIA | 0.82 | 0.79 | 0.8 | 0.73 | 0.81 | 0.68 | 0.7 | 0.84 |
| 8BTZ | 0.88 | 0.87 | 0.89 | 0.9 | 0.79 | 0.62 | 0.77 | 0.45 |
| 8BU8 | 0.86 | 0.87 | 0.83 | 0.83 | 0.71 | 0.58 | 0.11 | 0.66 |
| 8BWT | 0.91 | 0.89 | 0.89 | 0.92 | 0.83 | 0.87 | 0.82 | 0.79 |
| 8CLR | 0.97 | 0.97 | 0.91 | 0.88 | 0.94 | 0.83 | 0.94 | 0.76 |
| 8CQ1 | 0.9 | 0.9 | 0.87 | 0.88 | 0.84 | 0.81 | 0.72 | 0.62 |
| 8FB3 | 0.71 | 0.68 | 0.46 | 0.73 | 0.78 | 0.62 | 0.66 | 0.55 |
| 8FCS | 0.88 | 0.85 | 0.88 | 0.88 | 0.88 | 0.91 | 0.78 | 0.8 |
| 8FZA | 0.67 | 0.37 | 0.62 | 0.69 | 0.65 | 0.48 | 0.48 | 0.41 |
| 8HB8 | 0.61 | 0.59 | 0.56 | 0.5 | 0.53 | 0.48 | 0.42 | 0.2 |
| 8I43 | 0.81 | 0.78 | 0.81 | 0.85 | 0.69 | 0.67 | 0.74 | 0.69 |

| | | | | | | | | |
|---|---|---|---|---|---|---|---|---|
| **8I44** | 0.72 | 0.81 | 0.74 | 0.86 | 0.69 | 0.76 | 0.71 | 0.22 |
| **8I45** | 0.89 | 0.83 | 0.8 | 0.87 | 0.72 | 0.76 | 0.84 | 0.32 |
| **8I46** | 0.9 | 0.9 | 0.85 | 0.94 | 0.74 | 0.65 | 0.82 | 0.72 |
| **8ITS** | 0.84 | 0.85 | 0.84 | 0.87 | 0.88 | 0.82 | 0.81 | 0.45 |
| **8JHP** | 0.63 | 0.65 | 0.77 | 0.72 | 0.78 | 0.55 | 0.64 | 0.47 |
| **8K2Z** | 0.9 | 0.63 | 0.76 | 0.86 | 0.84 | 0.66 | 0.79 | 0.2 |
| **8K30** | 0.4 | 0.24 | 0.33 | 0.35 | 0.38 | 0.33 | 0.38 | 0.3 |
| **8K8B** | 0.66 | 0.78 | 0.71 | 0.74 | 0.66 | 0.72 | 0.72 | 0.34 |
| **8KEB** | 0.37 | 0.37 | 0.39 | 0.37 | 0.34 | 0.38 | 0.31 | 0.28 |
| **8Q4O** | 0.25 | 0.31 | 0.34 | 0.3 | 0.23 | 0.29 | 0.15 | 0.29 |
| **8Q6N** | 0.84 | 0.87 | 0.76 | 0.83 | 0.85 | 0.75 | 0.69 | 0.33 |
| **8QO2** | 0.57 | 0.55 | 0.55 | 0.58 | 0.66 | 0.52 | 0.52 | 0.58 |
| **8QO3** | 0.75 | 0.76 | 0.76 | 0.78 | 0.8 | 0.71 | 0.74 | 0.56 |
| **8QO4** | 0.7 | 0.67 | 0.67 | 0.68 | 0.71 | 0.68 | 0.66 | 0.62 |
| **8QO5** | 0.69 | 0.64 | 0.62 | 0.64 | 0.68 | 0.6 | 0.59 | 0.56 |
| **8SCF** | 0.82 | 0.82 | 0.81 | 0.8 | 0.81 | 0.76 | 0.79 | 0.3 |
| **8SCH** | 0.82 | 0.83 | 0.72 | 0.78 | 0.76 | 0.85 | 0.78 | 0.65 |
| **8T5O** | 0.41 | 0.41 | 0.44 | 0.39 | 0.47 | 0.4 | 0.27 | 0.31 |
| **8TNS** | 0.0 | 0.0 | 0.0 | 0.0 | 0.0 | 0.0 | 0.0 | 0.0 |
| **8TVZ** | 0.87 | 0.82 | 0.82 | 0.8 | 0.69 | 0.63 | 0.0 | 0.68 |
| **8UPT** | 0.77 | 0.74 | 0.69 | 0.71 | 0.72 | 0.63 | 0.68 | 0.57 |
| **8UTG** | 0.26 | 0.43 | 0.35 | 0.07 | 0.31 | 0.25 | 0.3 | 0.37 |
| **8UYG** | 0.86 | 0.79 | 0.57 | 0.77 | 0.76 | 0.69 | 0.66 | 0.54 |
| **8UYM** | 0.84 | 0.78 | 0.78 | 0.79 | 0.79 | 0.72 | 0.74 | 0.7 |
| **8UYP** | 0.89 | 0.74 | 0.77 | 0.83 | 0.72 | 0.72 | 0.75 | 0.59 |
| **8V1H** | 0.35 | 0.33 | 0.33 | 0.34 | 0.37 | 0.24 | 0.33 | 0.32 |
| **8V1I** | 0.36 | 0.33 | 0.4 | 0.33 | 0.32 | 0.32 | 0.35 | 0.24 |
| **8VCI** | 0.83 | 0.81 | 0.8 | 0.76 | 0.69 | 0.71 | 0.73 | 0.6 |
| **8VT5** | 0.87 | 0.87 | 0.76 | 0.88 | 0.82 | 0.74 | 0.81 | 0.58 |
| **8XZL** | 0.82 | 0.83 | 0.79 | 0.71 | 0.75 | 0.66 | 0.73 | 0.32 |
| **8ZAU** | 0.72 | 0.74 | 0.75 | 0.7 | 0.69 | 0.64 | 0.74 | 0.57 |
| **9BLM** | 0.83 | 0.89 | 0.62 | 0.81 | 0.86 | 0.8 | 0.68 | 0.43 |

| | | | | | | | | |
|---|---|---|---|---|---|---|---|---|
| **9BUN** | 0.74 | 0.76 | 0.67 | 0.71 | 0.69 | 0.69 | 0.61 | 0.47 |
| **9DE6** | 0.89 | 0.38 | 0.88 | 0.92 | 0.88 | 0.75 | 0.87 | 0.69 |
| **9DE7** | 0.93 | 0.88 | 0.78 | 0.88 | 0.86 | 0.77 | 0.84 | 0.56 |
| **9ECQ** | 0.98 | 0.92 | 0.8 | 0.92 | 0.87 | 0.89 | 0.78 | 0.78 |
| **9ECR** | 0.86 | 0.73 | 0.74 | 0.85 | 0.82 | 0.75 | 0.67 | 0.56 |
| **9EOW** | 0.8 | 0.85 | 0.84 | 0.87 | 0.78 | 0.77 | 0.75 | 0.35 |
| **9EWW** | 0.74 | 0.68 | 0.47 | 0.64 | 0.64 | 0.49 | 0.26 | 0.53 |
| **9FM4** | 0.79 | 0.78 | 0.73 | 0.82 | 0.8 | 0.8 | 0.8 | 0.56 |
| **9FO8** | 0.77 | 0.76 | 0.74 | 0.82 | 0.79 | 0.7 | 0.8 | 0.61 |
| **9FO9** | 0.77 | 0.76 | 0.8 | 0.78 | 0.77 | 0.67 | 0.75 | 0.74 |
| **9G7C** | 0.78 | 0.66 | 0.55 | 0.71 | 0.73 | 0.66 | 0.6 | 0.62 |
| **9IO0** | 0.92 | 0.9 | 0.83 | 0.95 | 0.74 | 0.72 | 0.79 | 0.81 |
| **9IO1** | 0.74 | 0.74 | 0.8 | 0.78 | 0.74 | 0.72 | 0.74 | 0.48 |
| **9IOR** | 0.82 | 0.82 | 0.76 | 0.78 | 0.59 | 0.71 | 0.73 | 0.66 |
| **9IOS** | 0.93 | 0.87 | 0.9 | 0.9 | 0.79 | 0.68 | 0.65 | 0.57 |
| **9IOU** | 0.85 | 0.8 | 0.76 | 0.84 | 0.65 | 0.66 | 0.79 | 0.69 |
| **R1205** | 0.6 | 0.57 | 0.44 | 0.59 | 0.57 | 0.58 | 0.46 | 0.48 |
| **R1209** | 0.78 | 0.8 | 0.76 | 0.79 | 0.76 | 0.79 | 0.76 | 0.72 |
| **R1211** | 0.88 | 0.88 | 0.86 | 0.89 | 0.84 | 0.76 | 0.77 | 0.61 |
| **R1212** | 0.22 | 0.26 | 0.28 | 0.21 | 0.34 | 0.28 | 0.22 | 0.29 |
| **R1248** | 0.39 | 0.38 | 0.45 | 0.35 | 0.46 | 0.39 | 0.32 | 0.36 |
| **R1263** | 0.86 | 0.92 | 0.75 | 0.84 | 0.86 | 0.96 | 0.86 | 0.78 |
| **R1271** | 0.82 | 0.82 | 0.79 | 0.82 | 0.81 | 0.79 | 0.72 | 0.36 |
| **R1288** | 0.68 | 0.68 | 0.64 | 0.6 | 0.65 | 0.64 | 0.65 | 0.38 |
| **R1293** | 0.68 | 0.76 | 0.71 | 0.76 | 0.69 | 0.66 | 0.6 | 0.5 |
| **R1296** | 0.86 | 0.83 | 0.83 | 0.86 | 0.84 | 0.82 | 0.74 | 0.62 |

**Table S4.** lDDT for the whole benchmarked dataset of single-chain RNA molecules across all tested deep learning methods for single-chain RNA prediction.

| PDB | AF3 | Boltz | Chai | HF3 | NuFold | RF2NA | RhoFold | trRosetta |
|---|---|---|---|---|---|---|---|---|
| **7EXY** | 0.48 | 0.11 | 0.62 | 0.53 | 0.52 | 0.11 | 0.6 | 0.27 |
| **7KUB** | 0.71 | 0.68 | 0.7 | 0.65 | 0.62 | 0.48 | 0.52 | 0.54 |
| **7KUC** | 0.78 | 0.64 | 0.75 | 0.58 | 0.8 | 0.61 | 0.61 | 0.73 |
| **7KUD** | 0.42 | 0.69 | 0.74 | 0.12 | 0.18 | 0.26 | 0.7 | 0.47 |
| **7MKT** | 0.26 | 0.04 | 0.25 | 0.0 | 0.25 | 0.05 | 0.19 | 0.18 |
| **7PS8** | 0.22 | 0.1 | 0.19 | 0.16 | 0.18 | 0.03 | 0.19 | 0.18 |
| **7Q48** | 0.25 | 0.14 | 0.23 | 0.18 | 0.18 | 0.09 | 0.22 | 0.22 |
| **7Q6L** | 0.26 | 0.22 | 0.28 | 0.2 | 0.23 | 0.15 | 0.23 | 0.28 |
| **7QA2** | 0.28 | 0.23 | 0.25 | 0.25 | 0.26 | 0.14 | 0.23 | 0.24 |
| **7RWR** | 0.54 | 0.54 | 0.52 | 0.43 | 0.48 | 0.44 | 0.52 | 0.57 |
| **7SHX** | 0.48 | 0.5 | 0.54 | 0.56 | 0.44 | 0.28 | 0.48 | 0.47 |
| **7SXP** | 0.24 | 0.12 | 0.23 | 0.0 | 0.21 | 0.08 | 0.22 | 0.19 |
| **7UGA** | 0.78 | 0.66 | 0.61 | 0.67 | 0.64 | 0.34 | 0.64 | 0.69 |
| **7UMC** | 0.63 | 0.71 | 0.46 | 0.57 | 0.53 | 0.41 | 0.63 | 0.55 |
| **7UMD** | 0.65 | 0.6 | 0.57 | 0.54 | 0.46 | 0.38 | 0.6 | 0.46 |
| **7UME** | 0.67 | 0.67 | 0.66 | 0.59 | 0.6 | 0.3 | 0.63 | 0.5 |
| **7V06** | 0.68 | 0.75 | 0.79 | 0.73 | 0.54 | 0.52 | 0.67 | 0.49 |
| **7WIA** | 0.75 | 0.73 | 0.64 | 0.55 | 0.6 | 0.51 | 0.6 | 0.69 |
| **8BTZ** | 0.74 | 0.63 | 0.77 | 0.76 | 0.64 | 0.35 | 0.66 | 0.54 |
| **8BU8** | 0.65 | 0.57 | 0.61 | 0.26 | 0.51 | 0.13 | 0.19 | 0.57 |
| **8BWT** | 0.73 | 0.74 | 0.74 | 0.69 | 0.58 | 0.6 | 0.64 | 0.62 |
| **8CLR** | 0.78 | 0.75 | 0.7 | 0.55 | 0.61 | 0.41 | 0.78 | 0.61 |
| **8CQ1** | 0.77 | 0.73 | 0.75 | 0.75 | 0.75 | 0.58 | 0.67 | 0.68 |
| **8FB3** | 0.63 | 0.42 | 0.25 | 0.29 | 0.78 | 0.1 | 0.69 | 0.48 |
| **8FCS** | 0.83 | 0.8 | 0.81 | 0.73 | 0.8 | 0.74 | 0.68 | 0.73 |
| **8FZA** | 0.66 | 0.19 | 0.41 | 0.55 | 0.33 | 0.08 | 0.53 | 0.55 |
| **8HB8** | 0.37 | 0.29 | 0.31 | 0.29 | 0.17 | 0.13 | 0.39 | 0.31 |
| **8I43** | 0.51 | 0.77 | 0.72 | 0.73 | 0.57 | 0.38 | 0.7 | 0.64 |

| | | | | | | | | |
|---|---|---|---|---|---|---|---|---|
| **8I44** | 0.69 | 0.8 | 0.71 | 0.61 | 0.6 | 0.5 | 0.69 | 0.52 |
| **8I45** | 0.68 | 0.58 | 0.66 | 0.7 | 0.38 | 0.15 | 0.69 | 0.52 |
| **8I46** | 0.71 | 0.74 | 0.71 | 0.69 | 0.47 | 0.13 | 0.68 | 0.64 |
| **8ITS** | 0.55 | 0.49 | 0.55 | 0.54 | 0.53 | 0.34 | 0.55 | 0.4 |
| **8JHP** | 0.49 | 0.5 | 0.63 | 0.47 | 0.43 | 0.4 | 0.5 | 0.5 |
| **8K2Z** | 0.68 | 0.4 | 0.56 | 0.63 | 0.43 | 0.16 | 0.61 | 0.49 |
| **8K30** | 0.72 | 0.4 | 0.59 | 0.6 | 0.54 | 0.25 | 0.65 | 0.68 |
| **8K8B** | 0.68 | 0.7 | 0.64 | 0.56 | 0.63 | 0.52 | 0.63 | 0.48 |
| **8KEB** | 0.79 | 0.56 | 0.54 | 0.74 | 0.53 | 0.25 | 0.64 | 0.49 |
| **8Q4O** | 0.27 | 0.03 | 0.22 | 0.22 | 0.26 | 0.09 | 0.21 | 0.22 |
| **8Q6N** | 0.65 | 0.64 | 0.62 | 0.57 | 0.77 | 0.47 | 0.58 | 0.54 |
| **8QO2** | 0.54 | 0.34 | 0.34 | 0.47 | 0.56 | 0.21 | 0.53 | 0.54 |
| **8QO3** | 0.7 | 0.68 | 0.66 | 0.68 | 0.65 | 0.3 | 0.66 | 0.58 |
| **8QO4** | 0.58 | 0.46 | 0.52 | 0.54 | 0.53 | 0.16 | 0.55 | 0.52 |
| **8QO5** | 0.67 | 0.48 | 0.47 | 0.53 | 0.6 | 0.21 | 0.55 | 0.48 |
| **8SCF** | 0.74 | 0.78 | 0.77 | 0.66 | 0.75 | 0.18 | 0.68 | 0.51 |
| **8SCH** | 0.73 | 0.64 | 0.62 | 0.57 | 0.62 | 0.42 | 0.6 | 0.57 |
| **8T5O** | 0.43 | 0.38 | 0.37 | 0.31 | 0.34 | 0.23 | 0.34 | 0.34 |
| **8TNS** | 0.23 | 0.28 | 0.28 | 0.02 | 0.23 | 0.03 | 0.22 | 0.23 |
| **8TVZ** | 0.62 | 0.44 | 0.58 | 0.58 | 0.41 | 0.22 | 0.0 | 0.43 |
| **8UPT** | 0.68 | 0.69 | 0.6 | 0.55 | 0.54 | 0.4 | 0.6 | 0.63 |
| **8UTG** | 0.29 | 0.16 | 0.23 | 0.02 | 0.21 | 0.12 | 0.21 | 0.21 |
| **8V1H** | 0.77 | 0.61 | 0.7 | 0.75 | 0.62 | 0.16 | 0.62 | 0.54 |
| **8V1I** | 0.72 | 0.42 | 0.63 | 0.53 | 0.5 | 0.32 | 0.6 | 0.62 |
| **8VCI** | 0.61 | 0.52 | 0.6 | 0.51 | 0.41 | 0.26 | 0.55 | 0.55 |
| **8VT5** | 0.69 | 0.56 | 0.45 | 0.62 | 0.43 | 0.27 | 0.67 | 0.67 |
| **8XZL** | 0.62 | 0.63 | 0.59 | 0.56 | 0.52 | 0.46 | 0.54 | 0.44 |
| **8ZAU** | 0.58 | 0.49 | 0.46 | 0.56 | 0.5 | 0.19 | 0.61 | 0.45 |
| **9BLM** | 0.79 | 0.78 | 0.5 | 0.56 | 0.73 | 0.57 | 0.58 | 0.43 |
| **9BUN** | 0.48 | 0.44 | 0.47 | 0.46 | 0.34 | 0.27 | 0.46 | 0.44 |
| **9DE6** | 0.73 | 0.21 | 0.8 | 0.77 | 0.76 | 0.52 | 0.64 | 0.65 |
| **9DE7** | 0.8 | 0.73 | 0.76 | 0.75 | 0.69 | 0.4 | 0.63 | 0.64 |

| | | | | | | | | |
|---|---|---|---|---|---|---|---|---|
| **9ECQ** | 0.75 | 0.76 | 0.78 | 0.68 | 0.76 | 0.3 | 0.68 | 0.71 |
| **9ECR** | 0.64 | 0.66 | 0.7 | 0.66 | 0.67 | 0.11 | 0.62 | 0.61 |
| **9EOW** | 0.7 | 0.56 | 0.71 | 0.66 | 0.74 | 0.22 | 0.72 | 0.56 |
| **9EWW** | 0.46 | 0.39 | 0.22 | 0.36 | 0.36 | 0.19 | 0.0 | 0.31 |
| **9FM4** | 0.67 | 0.64 | 0.68 | 0.56 | 0.64 | 0.42 | 0.64 | 0.63 |
| **9FO8** | 0.59 | 0.63 | 0.62 | 0.61 | 0.6 | 0.42 | 0.64 | 0.65 |
| **9FO9** | 0.6 | 0.63 | 0.66 | 0.58 | 0.69 | 0.53 | 0.66 | 0.68 |
| **9G7C** | 0.55 | 0.33 | 0.31 | 0.42 | 0.41 | 0.18 | 0.42 | 0.46 |
| **9IO0** | 0.8 | 0.81 | 0.75 | 0.67 | 0.53 | 0.52 | 0.7 | 0.72 |
| **9IO1** | 0.68 | 0.75 | 0.67 | 0.44 | 0.54 | 0.4 | 0.66 | 0.66 |
| **9IOR** | 0.76 | 0.8 | 0.7 | 0.75 | 0.52 | 0.52 | 0.68 | 0.7 |
| **9IOS** | 0.7 | 0.74 | 0.77 | 0.68 | 0.51 | 0.22 | 0.6 | 0.66 |
| **9IOU** | 0.71 | 0.77 | 0.64 | 0.55 | 0.58 | 0.42 | 0.68 | 0.67 |
| **R1205** | 0.41 | 0.37 | 0.28 | 0.45 | 0.24 | 0.3 | 0.33 | 0.34 |
| **R1209** | 0.6 | 0.47 | 0.58 | 0.57 | 0.5 | 0.35 | 0.54 | 0.54 |
| **R1211** | 0.63 | 0.59 | 0.62 | 0.73 | 0.59 | 0.26 | 0.53 | 0.53 |
| **R1212** | 0.39 | 0.42 | 0.5 | 0.39 | 0.45 | 0.03 | 0.41 | 0.42 |
| **R1248** | 0.18 | 0.0 | 0.13 | 0.15 | 0.16 | 0.07 | 0.15 | 0.17 |
| **R1263** | 0.63 | 0.55 | 0.56 | 0.58 | 0.6 | 0.83 | 0.62 | 0.75 |
| **R1271** | 0.73 | 0.57 | 0.73 | 0.72 | 0.48 | 0.28 | 0.57 | 0.52 |
| **R1288** | 0.64 | 0.62 | 0.54 | 0.41 | 0.45 | 0.48 | 0.55 | 0.39 |
| **R1293** | 0.55 | 0.48 | 0.54 | 0.6 | 0.3 | 0.2 | 0.46 | 0.51 |
| **R1296** | 0.69 | 0.57 | 0.58 | 0.69 | 0.58 | 0.36 | 0.55 | 0.55 |

**Table S5.** Mean scores and success rates for single-chain RNA prediction, shown per method.

| | **AlphaFold3** | **Boltz-1** | **RF2NA** | **Chai1** | **RhoFold+** | **NuFold** | **HelixFold3** | **trRosettaRNA** |
|---|---|---|---|---|---|---|---|---|
| **TM-score** | 0.326/19% | 0.326/14% | 0.296/10% | 0.291/12% | 0.276/12% | 0.325/19% | 0.287/8% | 0.252/8% |
| **INF-ALL** | 0.709/59% | 0.693/57% | 0.637/33% | 0.676/53% | 0.623/36% | 0.670/44% | 0.696/58% | 0.511/13% |
| **lDDT** | 0.599/15% | 0.526/14% | 0.307/1% | 0.557/13% | 0.531/1% | 0.504/9% | 0.520/7% | 0.508/1% |
| **GDT-TS** | 0.474/49% | 0.481/55% | 0.444/45% | 0.435/43% | 0.441/47% | 0.473/49% | 0.438/45% | 0.390/34% |

**Table S6.** DockQ scores for the whole benchmark dataset of RNA-RNA and RNA-Protein complexes across all tested deep learning methods for complex prediction.

| PDB | AF3 | Boltz-1 | HF3 | RF2NA |
|---|---|---|---|---|
| **7DA7** | 0,017 | 0,139 | 0,13 | 0,005 |
| **7ECJ** | 0,819 | 0,813 | 0,823 | 0,843 |
| **7ECK** | 0,742 | 0,776 | 0,804 | 0,853 |
| **7P0V** | 0,24 | 0,349 | 0,078 | 0,091 |
| **7PDU** | 0,22 | 0,231 | 0,213 | 0,102 |
| **7PMM** | 0,019 | 0,172 | 0,018 | 0,019 |
| **7QR3** | 0,208 | 0,229 | 0,052 | 0,238 |
| **7QR4** | 0,604 | 0,691 | 0,07 | 0,616 |
| **7SC6** | 0,218 | 0,252 | 0,252 | 0,258 |
| **7SCQ** | 0,215 | 0,257 | 0,229 | 0,251 |
| **7TNX** | 0,814 | 0,479 | 0,307 | 0,037 |
| **7TNY** | 0,76 | 0,673 | 0,3 | 0,23 |
| **7TO0** | 0,734 | 0,293 | 0,595 | 0,054 |
| **7TO1** | 0,22 | 0,177 | 0,029 | 0,041 |
| **7TO2** | 0,008 | 0,009 | 0,008 | 0,037 |
| **7U2A** | 0,7 | 0,586 | 0,326 | 0,272 |
| **7UIM** | 0,035 | 0,008 | 0,007 | 0,003 |
| **7UR5** | 0,009 | 0,014 | 0,114 | 0,028 |
| **7UU3** | 0,159 | 0,154 | 0,155 | 0,064 |
| **7V94** | 0,461 | 0,273 | 0,096 | 0,009 |
| **7XJZ** | 0,312 | 0,33 | 0,283 | 0,042 |
| **7YFQ** | 0,591 | 0,594 | 0,17 | 0,028 |
| **7YFX** | 0,18 | 0,313 | 0,048 | 0,009 |
| **7YFY** | 0,614 | 0,528 | 0,223 | 0,018 |
| **7YG6** | 0,402 | 0,419 | 0,368 | 0,023 |
| **7YGN** | 0,782 | 0,723 | 0,024 | 0,011 |
| **7YOJ** | 0,171 | 0,2 | 0,154 | 0,012 |
| **8AF0** | 0,196 | 0,136 | 0,134 | 0,024 |
| **8B9K** | 0,016 | 0,041 | 0,006 | 0,015 |
| **8CTI** | 0,165 | 0,012 | 0,012 | 0,015 |
| **8D9G** | 0,109 | 0,112 | 0,073 | 0,001 |
| **8DC2** | 0,259 | 0,178 | 0,179 | 0,006 |
| **8DMB** | 0,326 | 0,202 | 0,176 | 0,009 |

| | | | | |
|---|---|---|---|---|
| **8DVR** | 0,145 | 0,056 | 0,046 | 0,065 |
| **8DVS** | 0,19 | 0,145 | 0,033 | 0,025 |
| **8E28** | 0,048 | 0,381 | 0,039 | 0,45 |
| **8E29** | 0,381 | 0,376 | 0,351 | 0,283 |
| **8E2A** | 0,06 | 0,326 | 0,05 | 0,172 |
| **8FTI** | 0,008 | 0,009 | 0,013 | 0,013 |
| **8GZP** | 0,028 | 0,012 | 0,005 | 0,012 |
| **8GZR** | 0,124 | 0,152 | 0,02 | 0,008 |
| **8H1B** | 0,278 | 0,206 | 0,046 | 0,282 |
| **8H1J** | 0,14 | 0,251 | 0,101 | 0,006 |
| **8HB1** | 0,419 | 0,278 | 0,413 | 0,074 |
| **8HB3** | 0,154 | 0,2 | 0,378 | 0,075 |
| **8HBA** | 0,009 | 0,017 | 0,007 | 0,013 |
| **8HIO** | 0,018 | 0,038 | 0,025 | 0,017 |
| **8HKE** | 0,128 | 0,29 | 0,134 | 0,019 |
| **8HNT** | 0,124 | 0,008 | 0,015 | 0,009 |
| **8HNW** | 0,808 | 0,751 | 0,173 | 0,014 |
| **8HUD** | 0,441 | 0,698 | 0,174 | 0,016 |
| **8HZJ** | 0,044 | 0,04 | 0,012 | 0,017 |
| **8HZK** | 0,026 | 0,065 | 0,014 | 0,012 |
| **8HZL** | 0,011 | 0,012 | 0,008 | 0,003 |
| **8I3Q** | 0,009 | 0,043 | 0,015 | 0,009 |
| **8IBW** | 0,163 | 0,133 | 0,111 | 0,011 |
| **8IBY** | 0,005 | 0,016 | 0,022 | 0,033 |
| **8IBZ** | 0,006 | 0,015 | 0,007 | 0,009 |
| **8ID2** | 0,202 | 0,082 | 0,097 | 0,077 |
| **8IDF** | 0,141 | 0,16 | 0,013 | 0,013 |
| **8IEW** | 0,454 | 0,439 | 0,166 | 0,009 |
| **8IFK** | 0,37 | 0,663 | 0,102 | 0,006 |
| **8IN8** | 0,539 | 0,708 | 0,113 | 0,021 |
| **8IPM** | 0,498 | 0,477 | 0,018 | 0,024 |
| **8J8H** | 0,625 | 0,65 | 0,108 | 0,009 |
| **8J9G** | 0,016 | 0,018 | 0,018 | 0,013 |
| **8JL0** | 0,426 | 0,575 | 0,105 | 0,008 |
| **8JTJ** | 0,643 | 0,546 | 0,252 | 0,014 |
| **8JTR** | 0,307 | 0,184 | 0,268 | 0,014 |
| **8K0Y** | 0,007 | 0,234 | 0,011 | 0,026 |
| **8KAG** | 0,499 | 0,447 | 0,131 | 0,019 |
| **8OKD** | 0,579 | 0,136 | 0,006 | 0,153 |

| | | | | |
|---|---|---|---|---|
| **8OMR** | 0,741 | 0,281 | 0,737 | 0,282 |
| **8OPP** | 0,035 | 0,033 | 0,024 | 0,034 |
| **8PJB** | 0,249 | 0,107 | 0,108 | 0,164 |
| **8PM4** | 0,172 | 0,285 | 0,167 | 0,01 |
| **8PVV** | 0,244 | 0,189 | 0,079 | 0,014 |
| **8QG0** | 0,445 | 0,247 | 0,109 | 0,024 |
| **8R0S** | 0,602 | 0,607 | 0,232 | 0,115 |
| **8SCZ** | 0,645 | 0,437 | 0,023 | 0,023 |
| **8SD0** | 0,81 | 0,564 | 0,044 | 0,217 |
| **8SFH** | 0,426 | 0,421 | 0,113 | 0,035 |
| **8SFI** | 0,744 | 0,684 | 0,097 | 0,026 |
| **8SFJ** | 0,54 | 0,502 | 0,102 | 0,036 |
| **8SFL** | 0,455 | 0,428 | 0,096 | 0,025 |
| **8SFN** | 0,66 | 0,559 | 0,09 | 0,025 |
| **8SFR** | 0,748 | 0,604 | 0,098 | 0,028 |
| **8SH5** | 0,589 | 0,682 | 0,501 | 0,035 |
| **8SQU** | 0,576 | 0,694 | 0,122 | 0,012 |
| **8SXT** | 0,295 | 0,32 | 0,284 | 0,197 |
| **8T29** | 0,724 | 0,578 | 0,713 | 0,02 |
| **8T2P** | 0,003 | 0,005 | 0,003 | 0,149 |
| **8UZA** | 0,645 | 0,606 | 0,144 | 0,012 |
| **8VFS** | 0,664 | 0,631 | 0,172 | 0,01 |
| **8VIV** | 0,794 | 0,835 | 0,05 | 0,065 |
| **8VMB** | 0,609 | 0,54 | 0,628 | 0,02 |
| **8WFA** | 0,188 | 0,167 | 0,142 | 0,019 |
| **8WRR** | 0,192 | 0,427 | 0,134 | 0,011 |
| **8WRS** | 0,74 | 0,204 | 0,155 | 0,141 |
| **8WRT** | 0,012 | 0,041 | 0,031 | 0,018 |
| **8WRU** | 0,185 | 0,182 | 0,099 | 0,01 |
| **8WRV** | 0,184 | 0,18 | 0,158 | 0,138 |
| **8WS9** | 0,572 | 0,183 | 0,163 | 0,142 |
| **8X5V** | 0,764 | 0,7 | 0,177 | 0,013 |
| **8XCC** | 0,234 | 0,207 | 0,159 | 0,006 |
| **8XTP** | 0,006 | 0,004 | 0,008 | 0,003 |
| **8XYC** | 0,208 | 0,172 | 0,196 | 0,006 |
| **8Y03** | 0,46 | 0,396 | 0,039 | 0,099 |
| **8Y04** | 0,875 | 0,566 | 0,236 | 0,071 |
| **8Y0C** | 0,733 | 0,692 | 0,172 | 0,014 |
| **8Y0D** | 0,727 | 0,63 | 0,135 | 0,009 |

| | | | | |
|---|---|---|---|---|
| **8Y7Z** | 0,491 | 0,631 | 0,1 | 0,008 |
| **8Y9L** | 0,005 | 0,017 | 0,015 | 0,01 |
| **8Y9M** | 0,164 | 0,093 | 0,132 | 0,009 |
| **8YDC** | 0,462 | 0,524 | 0,455 | 0,215 |
| **8Z1F** | 0,013 | 0,298 | 0,015 | 0,019 |
| **8Z9K** | 0,03 | 0,203 | 0,018 | 0,068 |
| **8ZMI** | 0,226 | 0,243 | 0,253 | 0,251 |
| **8ZNJ** | 0,461 | 0,119 | 0,118 | 0,008 |
| **8ZTY** | 0,124 | 0,016 | 0,017 | 0,035 |
| **9B2K** | 0,376 | 0,369 | 0,125 | 0,007 |
| **9BXE** | 0,749 | 0,761 | 0,638 | 0,571 |
| **9C0I** | 0,327 | 0,418 | 0,02 | 0,053 |
| **9CER** | 0,015 | 0,008 | 0,017 | 0,017 |
| **9CEU** | 0,495 | 0,208 | 0,091 | 0,015 |
| **9CMP** | 0,435 | 0,568 | 0,235 | 0,043 |
| **9CPD** | 0,763 | 0,802 | 0,75 | 0,808 |
| **9CPJ** | 0,726 | 0,749 | 0,843 | 0,699 |
| **9DCF** | 0,026 | 0,052 | 0,02 | 0,024 |
| **9DIB** | 0,007 | 0,007 | 0,005 | 0,01 |
| **9DIG** | 0,007 | 0,113 | 0,008 | 0,007 |
| **9DII** | 0,007 | 0,015 | 0,007 | 0,009 |
| **9DTT** | 0,011 | 0,009 | 0,009 | 0,01 |
| **9E7G** | 0,282 | 0,624 | 0,26 | 0,013 |
| **9ENE** | 0,621 | 0,137 | 0,022 | 0,246 |
| **9FCV** | 0,009 | 0,055 | 0,025 | 0,021 |
| **9GBZ** | 0,191 | 0,184 | 0,235 | 0,007 |
| **9H82** | 0,01 | 0,055 | 0,008 | 0,011 |
| **9H83** | 0,156 | 0,116 | 0,067 | 0,118 |
| **9I8B** | 0,698 | 0,646 | 0,615 | 0,641 |
| **9IIY** | 0,527 | 0,53 | 0,208 | 0,017 |
| **9IIZ** | 0,216 | 0,508 | 0,213 | 0,024 |
| **9IJ0** | 0,51 | 0,637 | 0,24 | 0,066 |
| **9IJ1** | 0,235 | 0,204 | 0,225 | 0,012 |
| **9IJ2** | 0,505 | 0,486 | 0,425 | 0,017 |
| **9IJ3** | 0,598 | 0,66 | 0,244 | 0,027 |
| **9IJ4** | 0,62 | 0,618 | 0,244 | 0,022 |
| **9IJ5** | 0,617 | 0,624 | 0,481 | 0,028 |
| **9JSP** | 0,671 | 0,735 | 0,141 | 0,022 |
| **9KAD** | 0,077 | 0,082 | 0,081 | 0,082 |

| | | | | |
|---|---|---|---|---|
| **9KPH** | 0,004 | 0,005 | 0,006 | 0,003 |
| **9KPO** | 0,01 | 0,011 | 0,007 | 0,003 |
| **9L6Y** | 0,793 | 0,706 | 0,199 | 0,12 |
| **9MX3** | 0,514 | 0,262 | 0,259 | 0,25 |
| **9NVU** | 0,113 | 0,15 | 0,122 | 0,007 |
| **M1209** | 0,487 | 0,485 | 0,454 | 0,016 |
| **M1293** | 0,639 | 0,591 | 0,708 | 0,02 |
| **M1296** | 0,615 | 0,444 | 0,654 | 0,284 |

**Table S7.** US-Align TM-score for the whole benchmarked dataset of RNA-RNA and RNA-Protein complexes across all tested deep learning methods for complex prediction.

| PDB | AF3 | Boltz | HF3 | RF2NA |
|---|---|---|---|---|
| **7DA7** | 0,68238 | 0,57465 | 0,62719 | 0,07523 |
| **7ECJ** | 0,40557 | 0,38658 | 0,37352 | 0,37754 |
| **7ECK** | 0,36327 | 0,33774 | 0,34593 | 0,40694 |
| **7P0V** | 0,77768 | 0,82717 | 0,32433 | 0,77706 |
| **7PDU** | 0,17907 | 0,1871 | 0,18779 | 0,19043 |
| **7PMM** | 0,48528 | 0,49073 | 0,17282 | 0,44737 |
| **7QR3** | 0,39158 | 0,40516 | 0,25644 | 0,4191 |
| **7QR4** | 0,77069 | 0,85496 | 0,35405 | 0,75873 |
| **7SC6** | 0,77964 | 0,77917 | 0,78353 | 0,7763 |
| **7SCQ** | 0,77089 | 0,78851 | 0,80694 | 0,77661 |
| **7TNX** | 0,98362 | 0,95107 | 0,25868 | 0,75037 |
| **7TNY** | 0,98441 | 0,97381 | 0,2446 | 0,7521 |
| **7TO0** | 0,98748 | 0,94507 | 0,97961 | 0,76202 |
| **7TO1** | 0,97538 | 0,96086 | 0,24711 | 0,75843 |
| **7TO2** | 0,9207 | 0,92805 | 0,91718 | 0,78521 |
| **7U2A** | 0,98435 | 0,97064 | 0,95207 | 0,92681 |
| **7UIM** | 0,5168 | 0,4655 | 0,48037 | 0,42106 |
| **7UR5** | 0,33999 | 0,33997 | 0,27816 | 0,21843 |
| **7UU3** | 0,31505 | 0,52534 | 0,41843 | 0,324 |
| **7V94** | 0,90962 | 0,75409 | 0,26342 | 0,18965 |
| **7XJZ** | 0,88173 | 0,83494 | 0,87286 | 0,86782 |
| **7YFQ** | 0,94467 | 0,93084 | 0,94417 | 0,85695 |
| **7YFX** | 0,96902 | 0,91365 | 0,32918 | 0,89305 |
| **7YFY** | 0,9344 | 0,90134 | 0,21376 | 0,87786 |
| **7YG6** | 0,96636 | 0,93558 | 0,92061 | 0,90728 |
| **7YGN** | 0,96658 | 0,9208 | 0,2671 | 0,91899 |
| **7YOJ** | 0,55721 | 0,31427 | 0,17996 | 0,18112 |
| **8AF0** | 0,47963 | 0,44992 | 0,47859 | 0,41133 |
| **8B9K** | 0,77317 | 0,78386 | 0,23704 | 0,80669 |
| **8CTI** | 0,14623 | 0,12883 | 0,11586 | 0,12765 |
| **8D9G** | 0,78669 | 0,59792 | 0,21521 | 0,04273 |
| **8DC2** | 0,2267 | 0,23532 | 0,21237 | 0,20627 |

| | | | | |
|---|---|---|---|---|
| **8DMB** | 0,68894 | 0,55588 | 0,59474 | 0,43294 |
| **8DVR** | 0,97202 | 0,92154 | 0,33038 | 0,75148 |
| **8DVS** | 0,97258 | 0,95852 | 0,21688 | 0,77372 |
| **8E28** | 0,98095 | 0,96964 | 0,27198 | 0,97211 |
| **8E29** | 0,95959 | 0,95379 | 0,95899 | 0,92375 |
| **8E2A** | 0,73535 | 0,72989 | 0,23854 | 0,73072 |
| **8FTI** | 0,39743 | 0,31596 | 0,22863 | 0,13854 |
| **8GZP** | 0,88433 | 0,70251 | 0,23755 | 0,66439 |
| **8GZR** | 0,59603 | 0,53753 | 0,30916 | 0,07532 |
| **8H1B** | 0,91553 | 0,90471 | 0,36195 | 0,91236 |
| **8H1J** | 0,69409 | 0,73203 | 0,21971 | 0,68244 |
| **8HB1** | 0,26654 | 0,26121 | 0,21682 | 0,13788 |
| **8HB3** | 0,26476 | 0,19766 | 0,24865 | 0,15076 |
| **8HBA** | 0,20181 | 0,18175 | 0,2041 | 0,18426 |
| **8HIO** | 0,72656 | 0,71955 | 0,703 | 0,27637 |
| **8HKE** | 0,89922 | 0,9099 | 0,1954 | 0,25108 |
| **8HNT** | 0,76931 | 0,67264 | 0,76499 | 0,34801 |
| **8HNW** | 0,96207 | 0,95648 | 0,23096 | 0,5144 |
| **8HUD** | 0,94158 | 0,90837 | 0,20313 | 0,52965 |
| **8HZJ** | 0,25012 | 0,15663 | 0,21519 | 0,21989 |
| **8HZK** | 0,15646 | 0,22655 | 0,2359 | 0,2142 |
| **8HZL** | 0,21044 | 0,22514 | 0,15519 | 0,17922 |
| **8I3Q** | 0,89348 | 0,80123 | 0,79597 | 0,30555 |
| **8IBW** | 0,90644 | 0,86954 | 0,24262 | 0,41667 |
| **8IBY** | 0,69892 | 0,72561 | 0,71502 | 0,59132 |
| **8IBZ** | 0,71375 | 0,72918 | 0,21358 | 0,55067 |
| **8ID2** | 0,39234 | 0,40109 | 0,18109 | 0,40613 |
| **8IDF** | 0,8883 | 0,88179 | 0,24869 | 0,66293 |
| **8IEW** | 0,80364 | 0,73911 | 0,26905 | 0,18359 |
| **8IFK** | 0,9666 | 0,95494 | 0,23947 | 0,45867 |
| **8IN8** | 0,96901 | 0,96218 | 0,23583 | 0,57859 |
| **8IPM** | 0,94243 | 0,95224 | 0,29408 | 0,86989 |
| **8J8H** | 0,96568 | 0,96416 | 0,28226 | 0,56216 |
| **8J9G** | 0,92547 | 0,904 | 0,26941 | 0,50171 |
| **8JL0** | 0,96514 | 0,93952 | 0,22576 | 0,47953 |
| **8JTJ** | 0,93881 | 0,9013 | 0,88178 | 0,37751 |

| | | | | |
|---|---|---|---|---|
| **8JTR** | 0,91679 | 0,81789 | 0,90913 | 0,56184 |
| **8K0Y** | 0,40995 | 0,24211 | 0,37985 | 0,29716 |
| **8KAG** | 0,77924 | 0,73098 | 0,19862 | 0,33769 |
| **8OKD** | 0,93424 | 0,74392 | 0,16875 | 0,74798 |
| **8OMR** | 0,95391 | 0,89134 | 0,96433 | 0,86706 |
| **8OPP** | 0,91676 | 0,91338 | 0,25527 | 0,43416 |
| **8PJB** | 0,92669 | 0,96137 | 0,23901 | 0,81857 |
| **8PM4** | 0,89967 | 0,90732 | 0,23339 | 0,49274 |
| **8PVV** | 0,75219 | 0,64515 | 0,30346 | 0,34557 |
| **8QG0** | 0,88685 | 0,71653 | 0,26352 | 0,37786 |
| **8R0S** | 0,9418 | 0,94469 | 0,23761 | 0,85986 |
| **8SCZ** | 0,96132 | 0,92831 | 0,28326 | 0,76782 |
| **8SD0** | 0,98451 | 0,96255 | 0,23677 | 0,77847 |
| **8SFH** | 0,73886 | 0,7533 | 0,23776 | 0,50922 |
| **8SFI** | 0,81756 | 0,90454 | 0,21078 | 0,2851 |
| **8SFJ** | 0,91453 | 0,93111 | 0,25286 | 0,30795 |
| **8SFL** | 0,88308 | 0,87613 | 0,19488 | 0,28258 |
| **8SFN** | 0,97897 | 0,93795 | 0,2307 | 0,33218 |
| **8SFR** | 0,93807 | 0,9083 | 0,21848 | 0,35464 |
| **8SH5** | 0,796 | 0,91206 | 0,86445 | 0,42137 |
| **8SQU** | 0,95349 | 0,95189 | 0,23469 | 0,53367 |
| **8SXT** | 0,97329 | 0,96105 | 0,95187 | 0,87415 |
| **8T29** | 0,88649 | 0,85284 | 0,88686 | 0,41879 |
| **8T2P** | 0,27536 | 0,26496 | 0,30018 | 0,26251 |
| **8UZA** | 0,94706 | 0,9001 | 0,83514 | 0,32341 |
| **8VFS** | 0,56308 | 0,42473 | 0,26751 | 0,21285 |
| **8VIV** | 0,98677 | 0,98451 | 0,9007 | 0,91329 |
| **8VMB** | 0,80322 | 0,81183 | 0,83655 | 0,448 |
| **8WFA** | 0,64092 | 0,6142 | 0,58334 | 0,16776 |
| **8WRR** | 0,76056 | 0,59731 | 0,20658 | 0,1928 |
| **8WRS** | 0,78284 | 0,62674 | 0,29219 | 0,19219 |
| **8WRT** | 0,67088 | 0,65712 | 0,20077 | 0,34267 |
| **8WRU** | 0,36546 | 0,36247 | 0,21141 | 0,18721 |
| **8WRV** | 0,24591 | 0,23865 | 0,27196 | 0,2136 |
| **8WS9** | 0,87178 | 0,60265 | 0,22242 | 0,14617 |
| **8X5V** | 0,95384 | 0,93225 | 0,23206 | 0,66527 |

| | | | | |
|---|---|---|---|---|
| **8XCC** | 0,32835 | 0,36532 | 0,38843 | 0,17443 |
| **8XTP** | 0,31137 | 0,16753 | 0,18356 | 0,10159 |
| **8XYC** | 0,75621 | 0,34103 | 0,1973 | 0,19681 |
| **8Y03** | 0,98372 | 0,79085 | 0,20213 | 0,78604 |
| **8Y04** | 0,89483 | 0,87396 | 0,81693 | 0,846 |
| **8Y0C** | 0,97885 | 0,91162 | 0,23683 | 0,32485 |
| **8Y0D** | 0,87298 | 0,82815 | 0,26562 | 0,46993 |
| **8Y7Z** | 0,97079 | 0,96892 | 0,22446 | 0,58415 |
| **8Y9L** | 0,33144 | 0,25834 | 0,24522 | 0,22683 |
| **8Y9M** | 0,35869 | 0,22221 | 0,19136 | 0,18751 |
| **8YDC** | 0,35945 | 0,41154 | 0,37799 | 0,16162 |
| **8Z1F** | 0,90223 | 0,91534 | 0,90319 | 0,86902 |
| **8Z9K** | 0,21981 | 0,24911 | 0,21575 | 0,19706 |
| **8ZMI** | 0,76712 | 0,78734 | 0,81184 | 0,78836 |
| **8ZNJ** | 0,83708 | 0,70436 | 0,2751 | 0,36913 |
| **8ZTY** | 0,90509 | 0,77088 | 0,24299 | 0,21067 |
| **9B2K** | 0,62131 | 0,46813 | 0,51632 | 0,07643 |
| **9BXE** | 0,36054 | 0,36525 | 0,32187 | 0,22662 |
| **9C0I** | 0,72677 | 0,70673 | 0,21248 | 0,69273 |
| **9CER** | 0,29769 | 0,21999 | 0,2057 | 0,19335 |
| **9CEU** | 0,83352 | 0,84067 | 0,22311 | 0,20472 |
| **9CMP** | 0,83141 | 0,8796 | 0,83206 | 0,75011 |
| **9CPD** | 0,3723 | 0,41024 | 0,33386 | 0,44313 |
| **9CPJ** | 0,36602 | 0,43385 | 0,48665 | 0,43372 |
| **9DCF** | 0,51036 | 0,48265 | 0,2342 | 0,44771 |
| **9DIB** | 0,27955 | 0,19055 | 0,21449 | 0,17011 |
| **9DIG** | 0,20794 | 0,24915 | 0,21747 | 0,20767 |
| **9DII** | 0,22615 | 0,18507 | 0,19235 | 0,21609 |
| **9DTT** | 0,69908 | 0,70334 | 0,23827 | 0,67592 |
| **9E7G** | 0,79653 | 0,86392 | 0,81707 | 0,44714 |
| **9ENE** | 0,94547 | 0,75799 | 0,22573 | 0,83554 |
| **9FCV** | 0,94187 | 0,93031 | 0,24242 | 0,28248 |
| **9GBZ** | 0,73822 | 0,75835 | 0,71892 | 0,3614 |
| **9H82** | 0,08563 | 0,08929 | 0,08556 | 0,08492 |
| **9H83** | 0,21739 | 0,22869 | 0,1587 | 0,26549 |
| **9I8B** | 0,44226 | 0,39022 | 0,33738 | 0,35249 |

| | | | | |
|---|---|---|---|---|
| **9IIY** | 0,82046 | 0,84463 | 0,23278 | 0,76315 |
| **9IIZ** | 0,87398 | 0,8762 | 0,28758 | 0,84357 |
| **9IJ0** | 0,96531 | 0,89928 | 0,24559 | 0,89297 |
| **9IJ1** | 0,86269 | 0,9334 | 0,24692 | 0,79405 |
| **9IJ2** | 0,89722 | 0,8787 | 0,88688 | 0,84138 |
| **9IJ3** | 0,89747 | 0,90681 | 0,23174 | 0,80687 |
| **9IJ4** | 0,89837 | 0,94005 | 0,30536 | 0,82411 |
| **9IJ5** | 0,91378 | 0,91342 | 0,86361 | 0,83706 |
| **9JSP** | 0,9287 | 0,95196 | 0,31057 | 0,64381 |
| **9KAD** | 0,30271 | 0,33702 | 0,2411 | 0,38966 |
| **9KPH** | 0,20907 | 0,19558 | 0,22268 | 0,1403 |
| **9KPO** | 0,16545 | 0,15807 | 0,21169 | 0,12563 |
| **9L6Y** | 0,93578 | 0,90055 | 0,32427 | 0,75667 |
| **9MX3** | 0,9207 | 0,85612 | 0,87274 | 0,8873 |
| **9NVU** | 0,50552 | 0,51557 | 0,23458 | 0,20092 |
| **M1209** | 0,70825 | 0,69966 | 0,70289 | 0,41442 |
| **M1293** | 0,86699 | 0,85049 | 0,92418 | 0,42098 |
| **M1296** | 0,84565 | 0,8054 | 0,82882 | 0,86068 |

**Table S8.** Mean TM-scores, DockQ scores, DockQ success rates (SR) and interface DockQ (IF DockQ) scores for each RNA Complex prediction method. Mean IF DockQ excludes scores for protein-protein interface DockQ and is averaged across RNA-protein, RNA-RNA interface DockQ.

| | **AlphaFold3** | **Boltz-1** | **HelixFold3** | **RF2NA** |
|---|---|---|---|---|
| **TM-score** | 0.711 | 0.680 | 0.391 | 0.485 |
| **DockQ / SR [%]** | 0.390/54.4% | 0.357/54.4% | 0.166/24.7% | 0.062/12% |
| **IF DockQ** | 0.322 | 0.289 | 0.065 | 0.145 |

**Table S9.** Mean interface DockQ scores for RNA-protein and RNA-RNA interfaces.

| | **AlphaFold3** | **Boltz-1** | **HelixFold3** | **RF2NA** |
|---|---|---|---|---|
| **RNA-protein** | 0.305 | 0.261 | 0.088 | 0.045 |
| **RNA-RNA** | 0.384 | 0.393 | 0.350 | 0.136 |

**Table S10.** Mean ipTM scores for each complex type across all methods.

| Complex type | AF3 | Boltz-1 | HF3 |
|---|---|---|---|
| **RNA-RNA** | 0.246 | 0.247 | 0.308 |
| **RNA-protein** | 0.585 | 0.523 | 0.362 |
| **RNA-DNA** | 0.410 | 0.399 | 0.585 |
| **RNA-protein/DNA** | 0.645 | 0.625 | 0.164 |

**Table S11.** Mean chain ipTM scores for each chain type for AlphaFold3 predictions.

| Chain type | N | Mean chain ipTM |
|---|---|---|
| **RNA** | 219 | 0.412 |
| **Protein** | 171 | 0.556 |
| **DNA** | 76 | 0.424 |

### Supplementary Figures

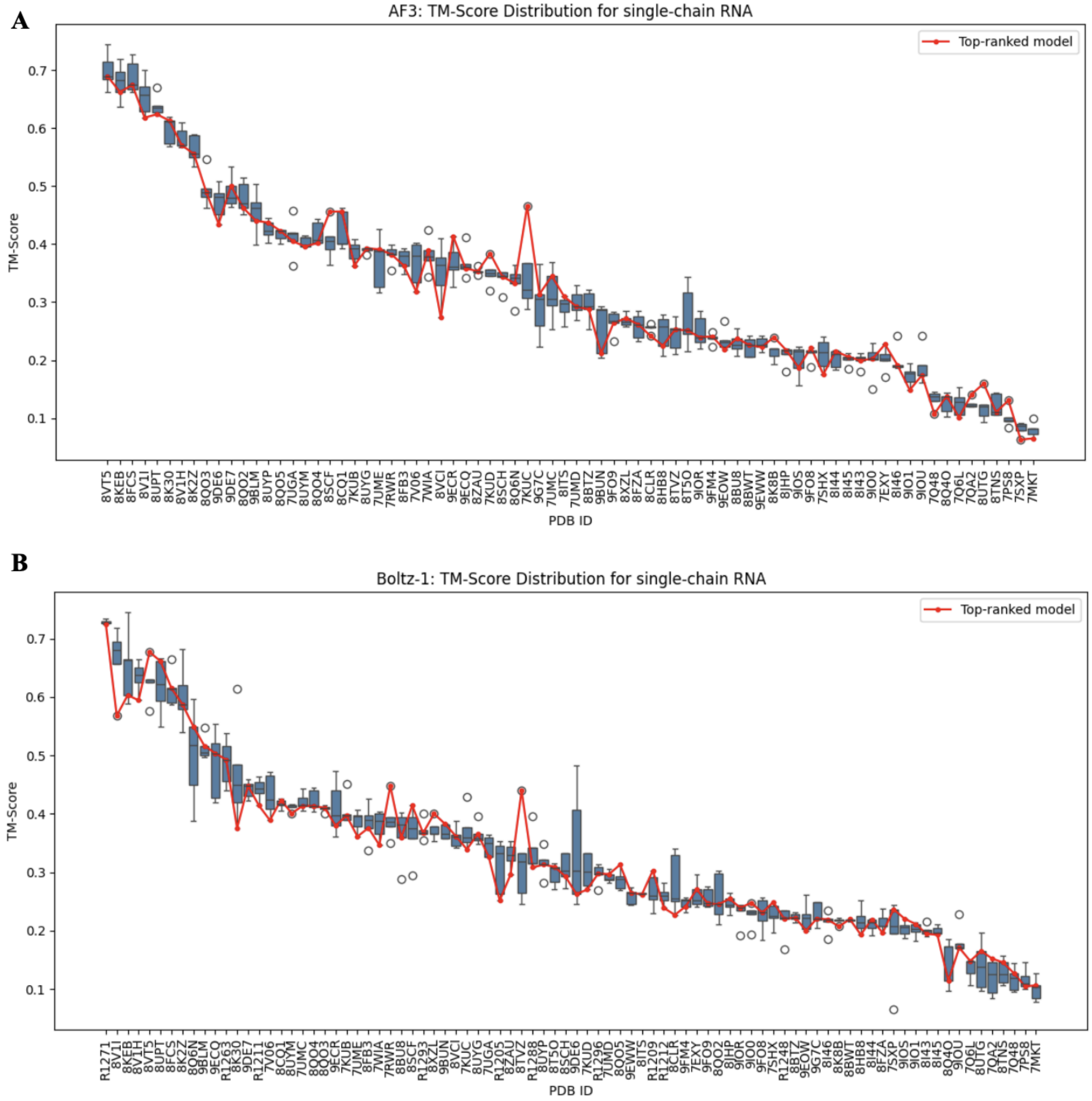


**Figure S1.** Distribution of TM-scores across $n$ = 5 generated single-chain RNA models for AlphaFold3 (A) and Boltz-1 (B). TM scores for each ID are shown as box plots, ordered from highest to lowest mean TM score. The red line across the plot connects the top-ranked models of each ID.

**A**

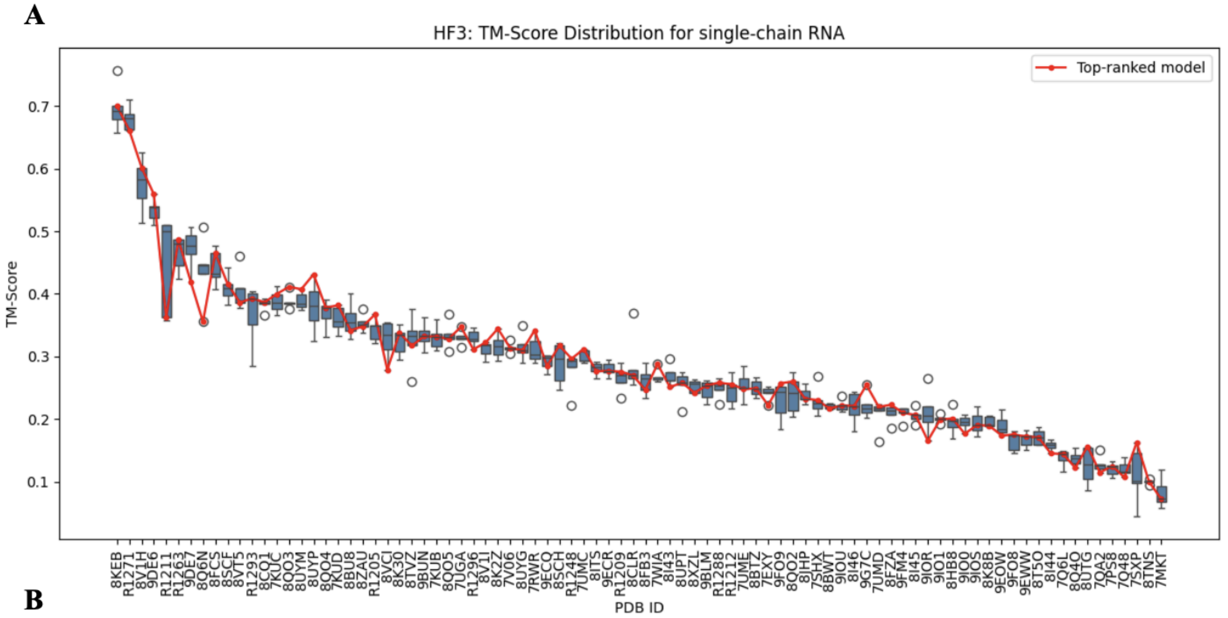


**B**

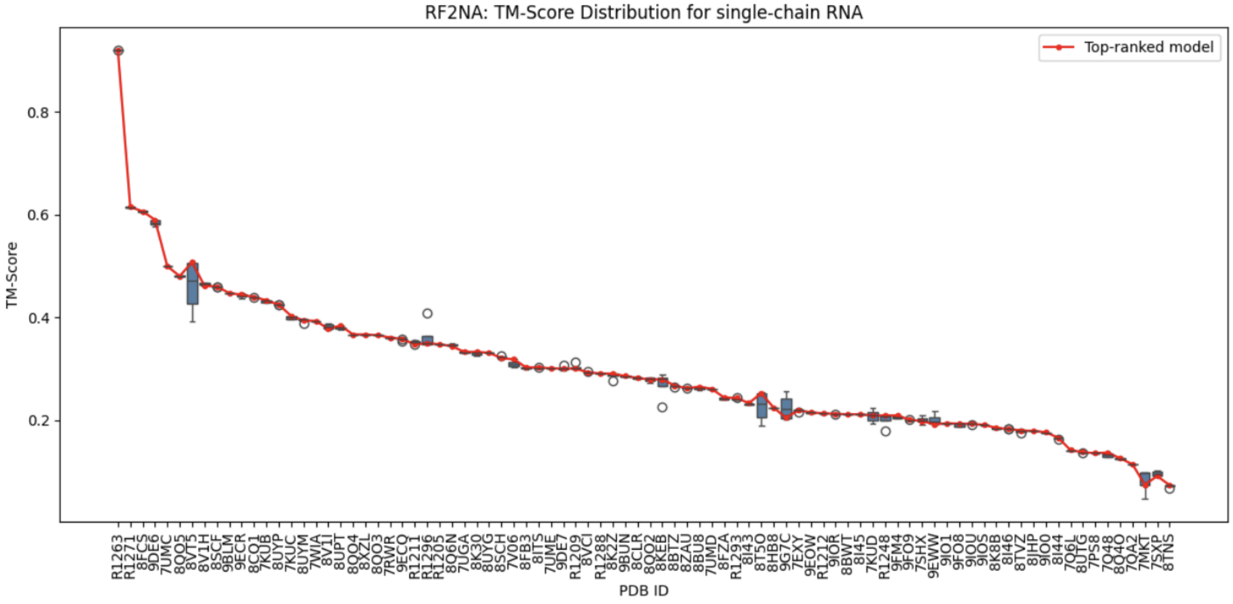


**Figure S2.** Distribution of TM-scores across *n = 5* generated single-chain RNA models for HF3 (A) and RF2NA1 (B). For each ID, TM scores are visualized as box plots, ordered from highest to lowest mean TM score. A red line connects the top-ranked model for each ID.

**A**

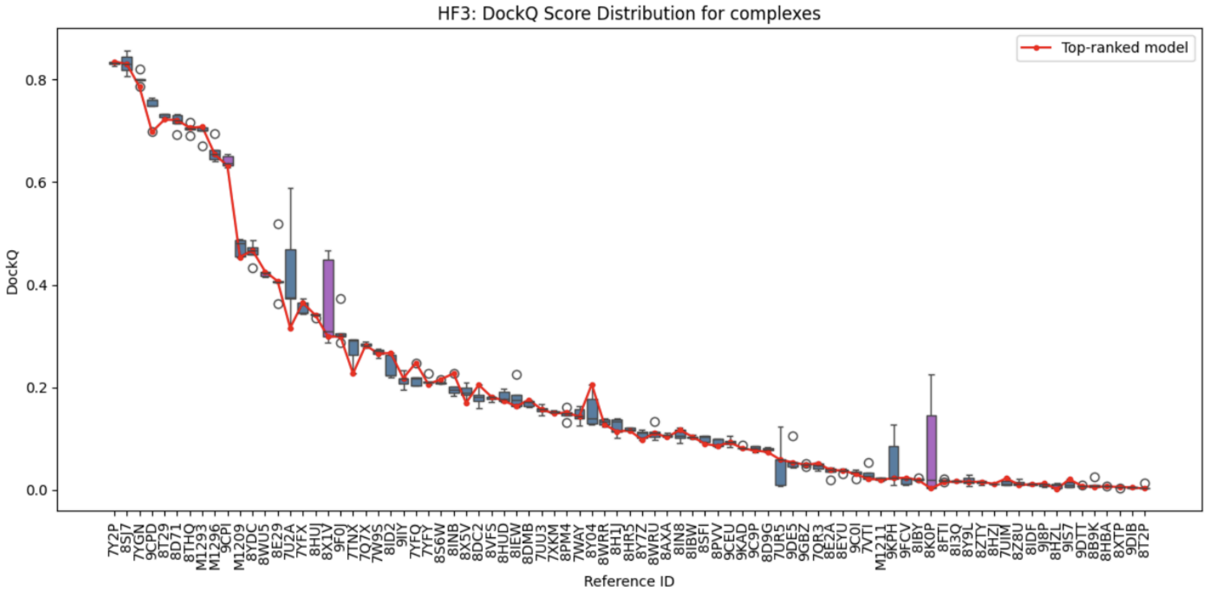


**B**

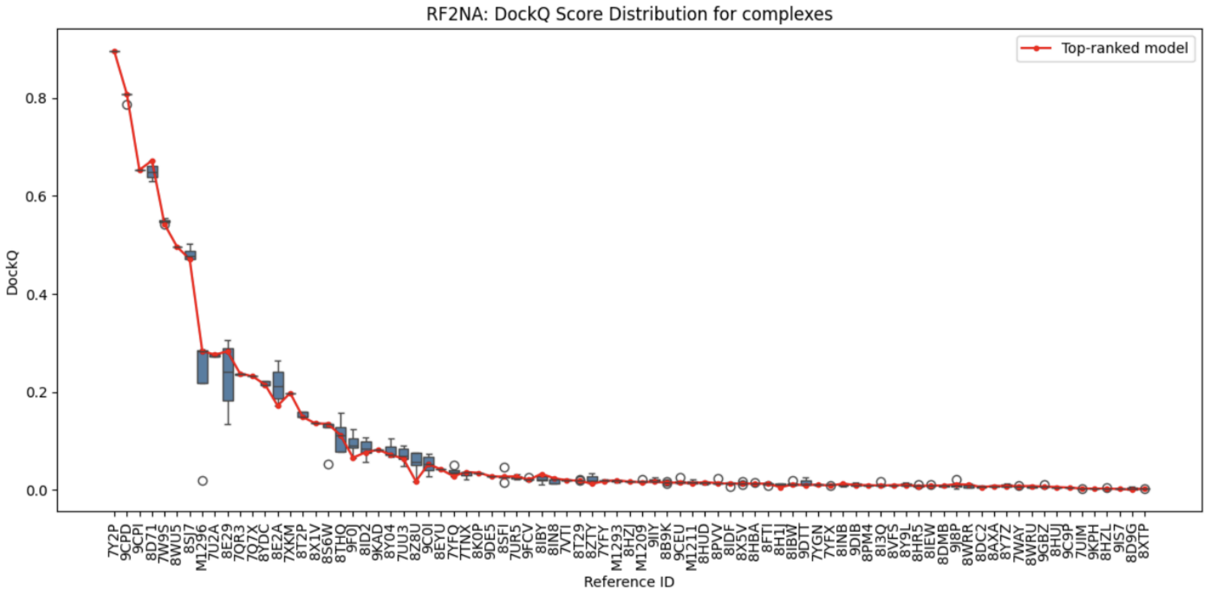


**Figure S3.** Distribution of DockQ scores across $n = 5$ generated RNA complex models for HF3 (A) and RF2NA1 (B). For each ID, DockQ scores are visualized as box plots, ordered from highest to lowest mean DockQ score. Box plots are colored blue for RNA-protein complexes and purple for RNA-RNA complexes. A red line connects the top-ranked model for each ID.

**A**

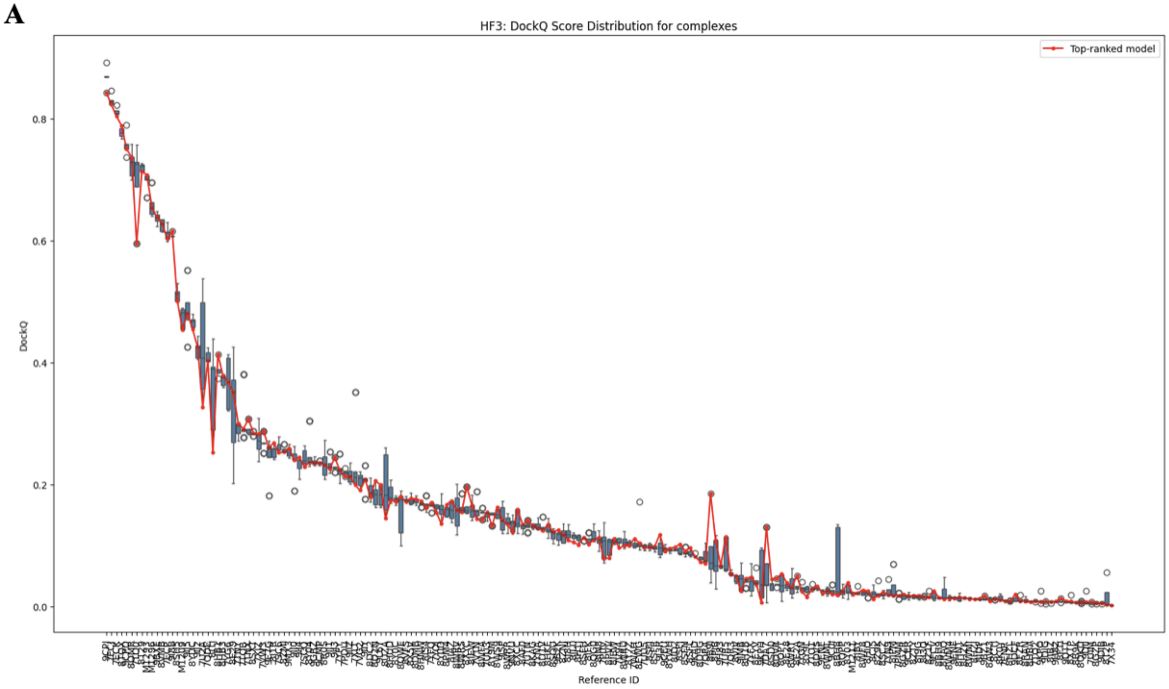


**B**

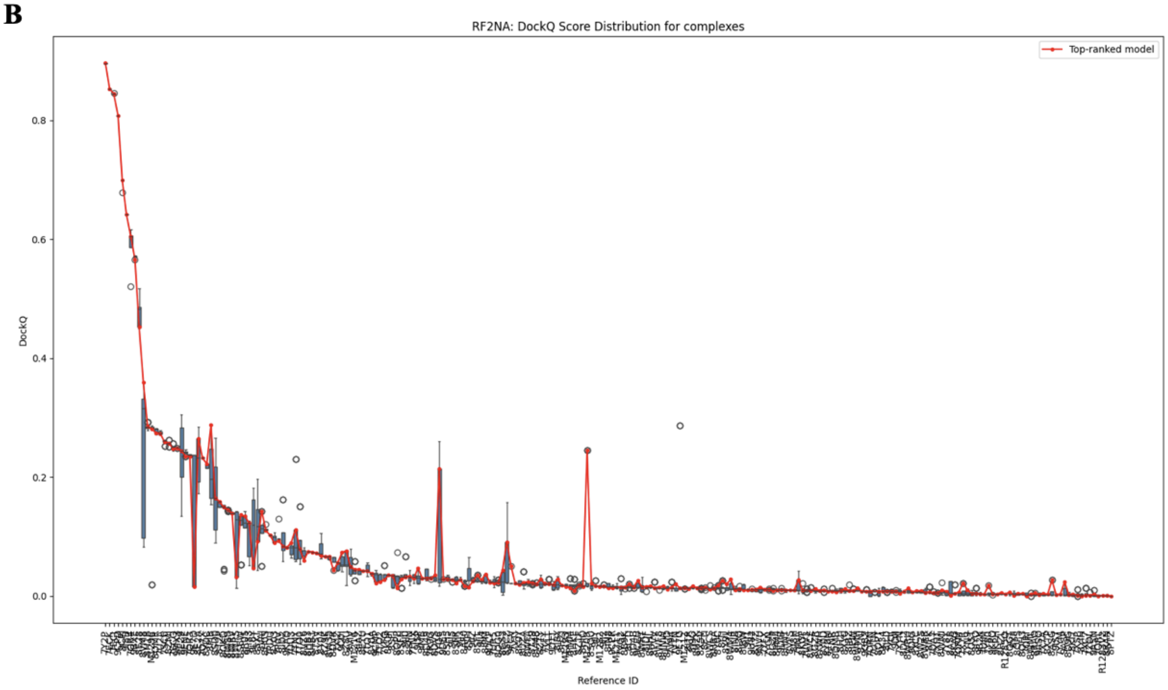


**Figure S4**. Distribution of DockQ scores across $n = 5$ generated RNA complex models for HF3 (A) and RF2NA1 (B). For each ID, DockQ scores are visualized as box plots, ordered by decreasing mean DockQ score. Box plots are colored blue for RNA-protein complexes and purple for RNA-RNA complexes. A red line connects the top-ranked model for each ID.

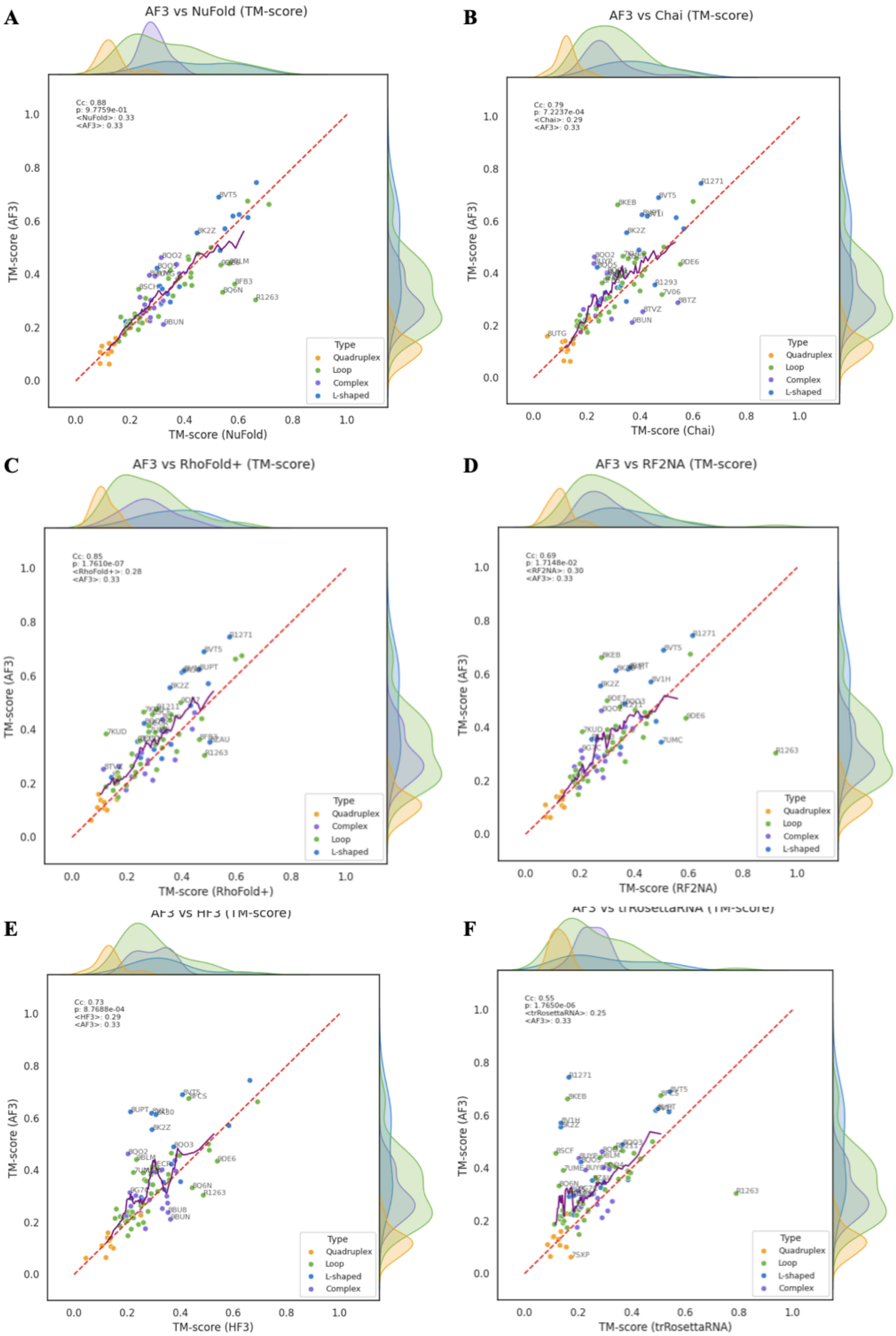

**Figure S5.** Pair plots showing TM-score correlation between the top-ranked method (AlphaFold3) against TM-scores of other methods. Cc denotes the Pearson correlation coefficient, paired T-test p-value by *p*.

A B

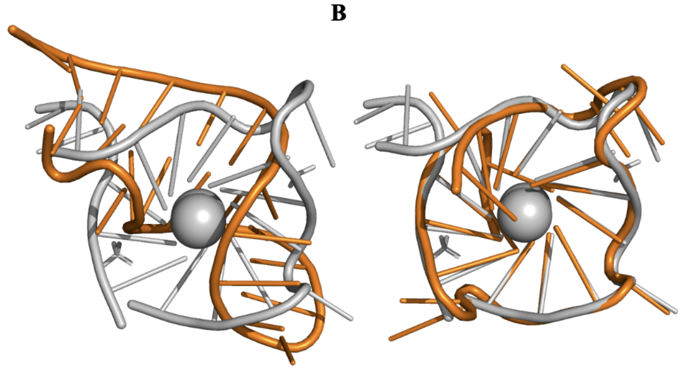

| | **8VT5** (AF3) | **8FCS** (Boltz-1) |
|---|---|---|
| TM-score | 0.063 | 0.237 |
| INF | 0.29 | 0.42 |
| GDT-TS | 0.34 | 0.79 |
| lDDT | 0.26 | 0.2 |
| RMSD | 10.528 | 0.796 |

**Figure S6.** Comparison of modelling 8VT5 between AlphaFold 3 (left) and Boltz-1 (right) where Boltz-1 predicted the structure with better accuracy, predicting the correct overall fold, while AlphaFold 3 failed.

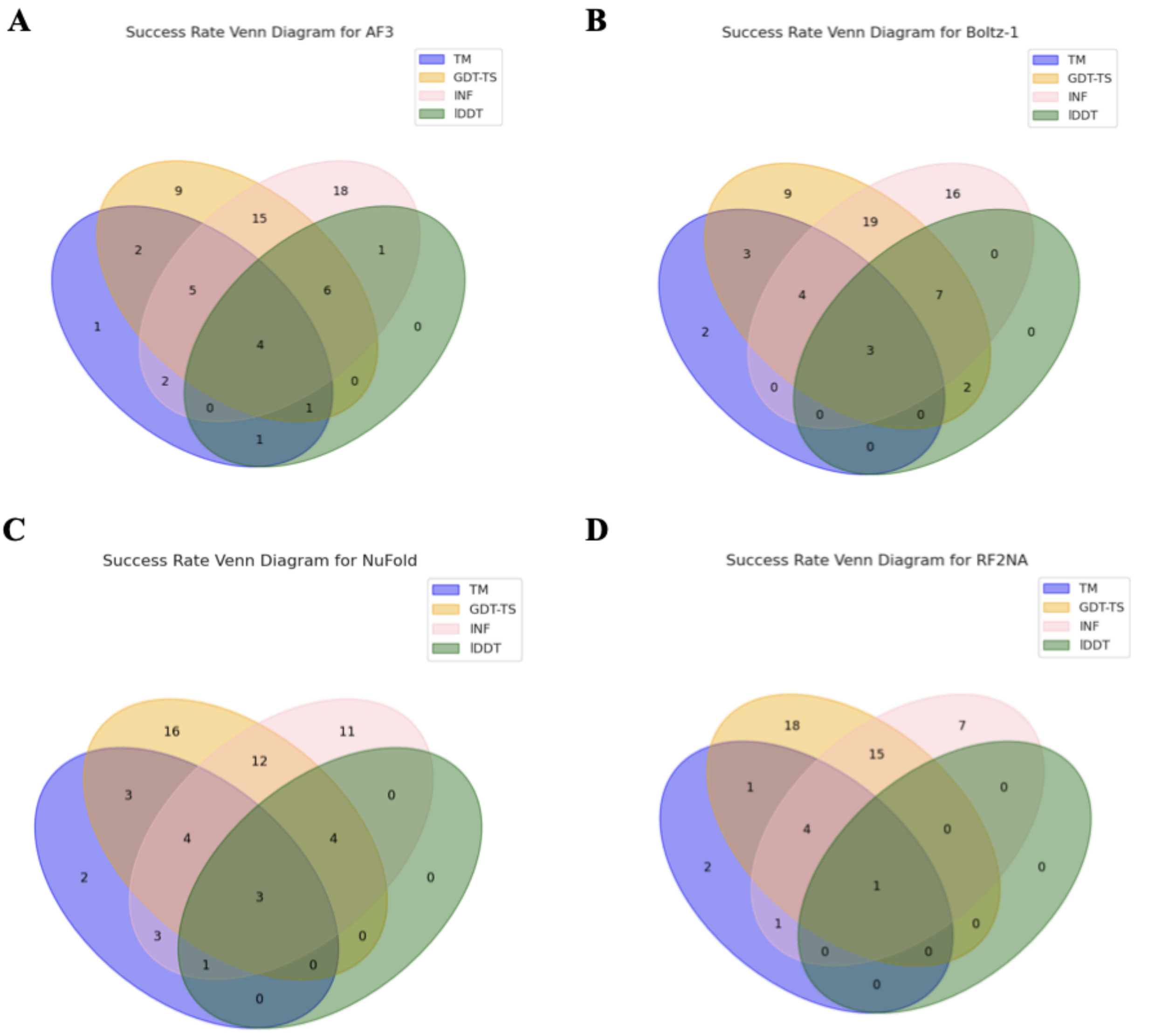


**Figure S7.** Venn diagrams showing the number of successful models based on the thresholds of different evaluation metrics, for (A) AlphaFold3, (B) Boltz-1, (C) NuFold and (D) RosettaFold2NA

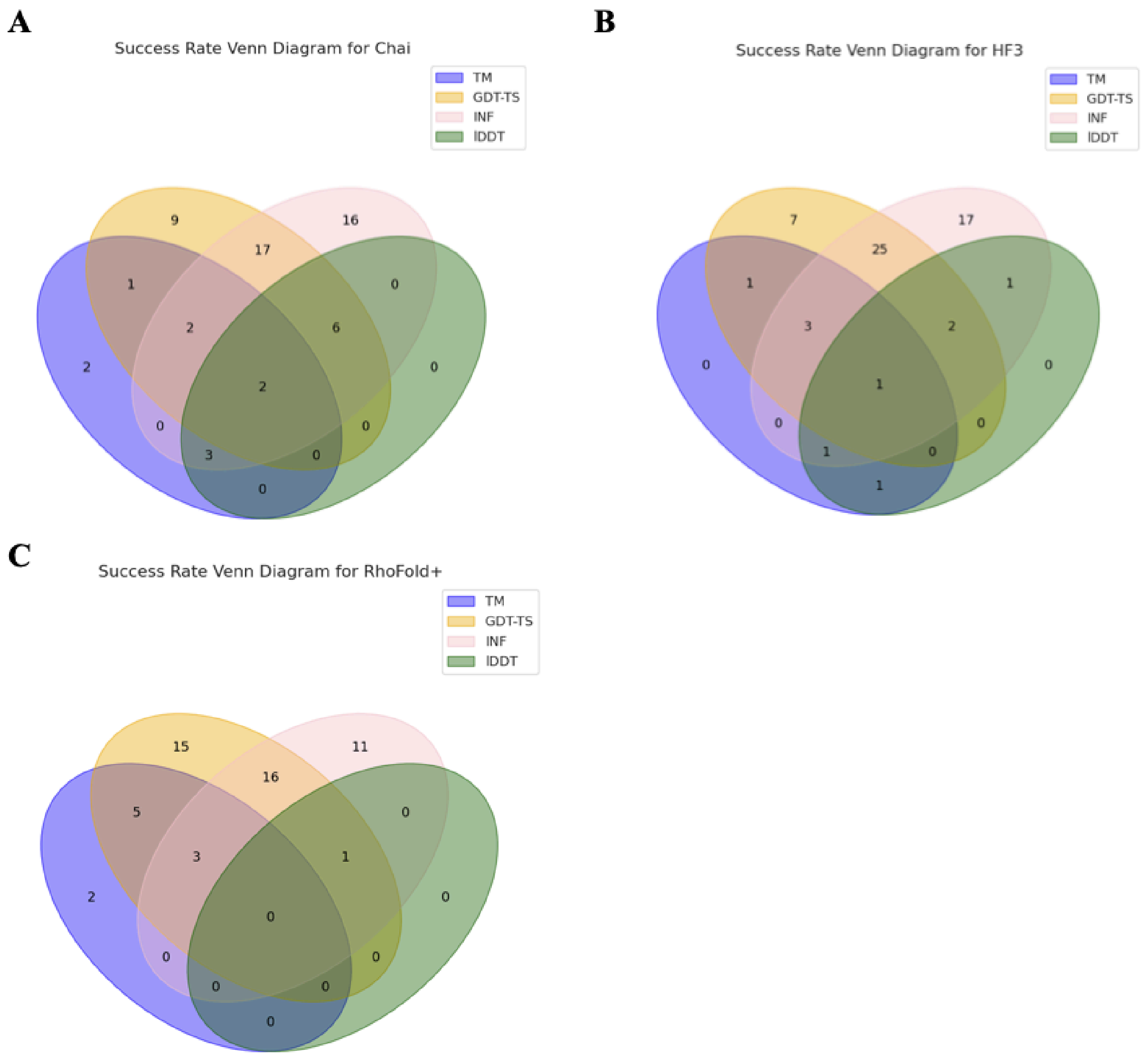


**Figure S8.** Venn diagrams showing the number of successful models based on the thresholds of different evaluation metrics, for (A) Chai-1, (B) HelixFold 3, (C) RhoFold+

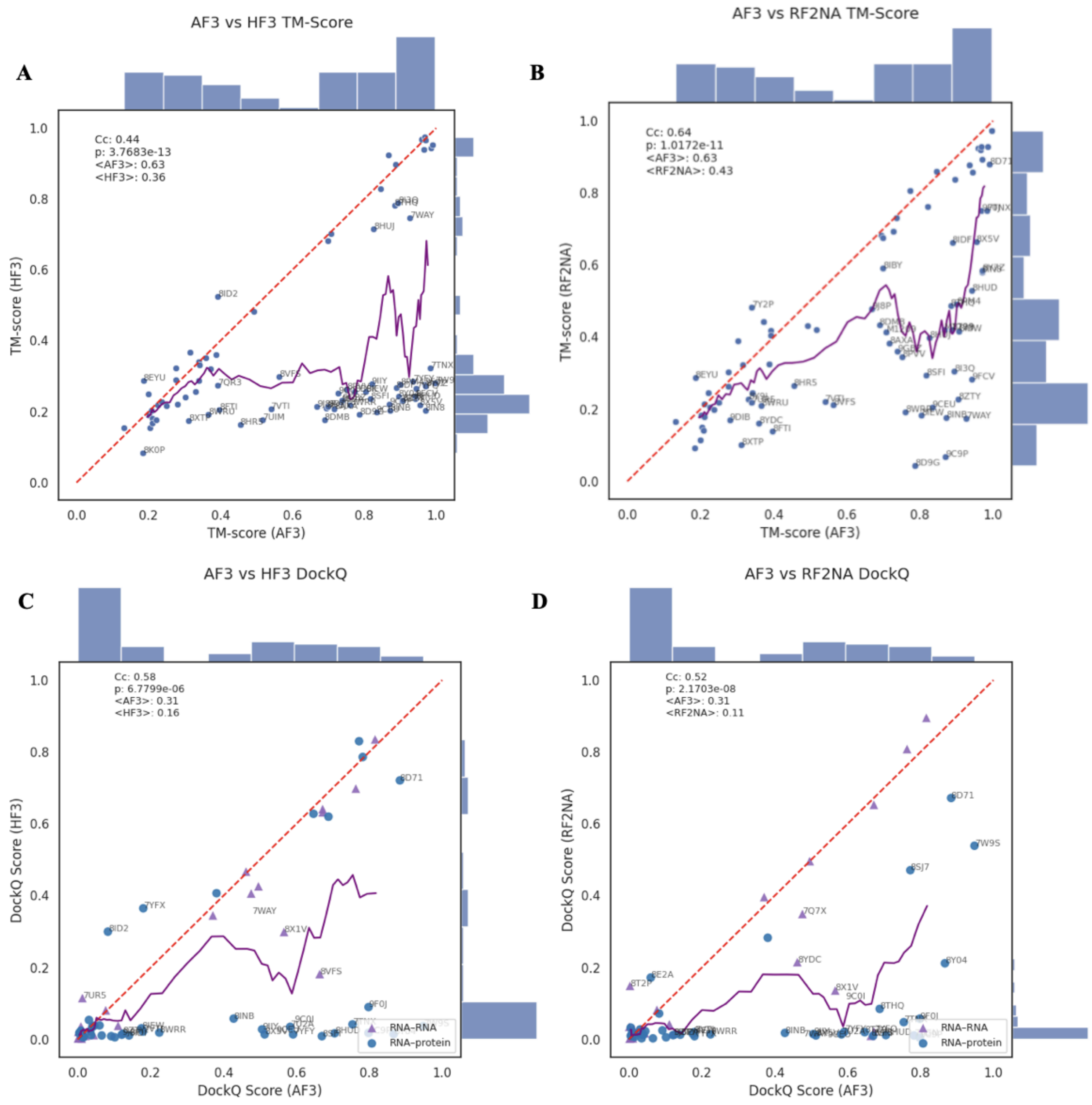


**Figure S9.** Scatter plots showing (A, B) TM-score and (C, D) Interface DockQ score correlation between the top-ranked method (AlphaFold3) against the corresponding scores of other methods. Cc denotes the Pearson correlation coefficient, and the paired T-test p-value by p. Significant differences in TM-scores and interface DockQ scores were found between AlphaFold3 and Boltz-1, HF3, and RF2NA, for both TM-score and interface DockQ score.

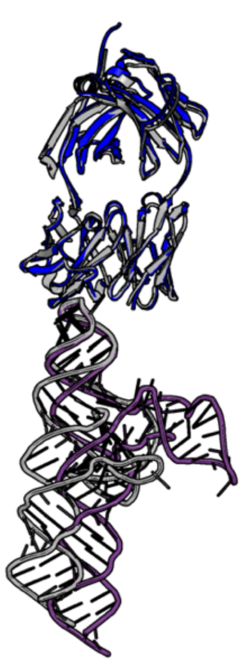

**8T29 (HF3)**
**RNA-protein DockQ 0.670**
RNA-RNA DockQ /
Total DockQ 0.713
TM-score 0.887

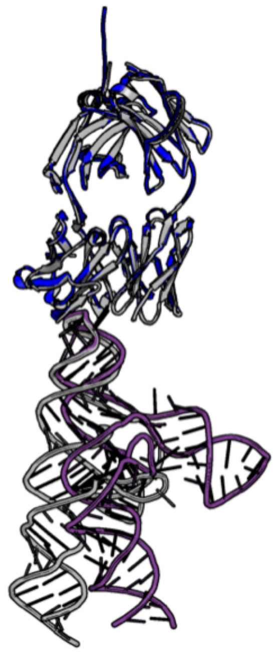

**8T29 (AF3)**
**RNA-protein DockQ 0.647**
RNA-RNA DockQ /
Total DockQ 0.724
TM-score 0.886

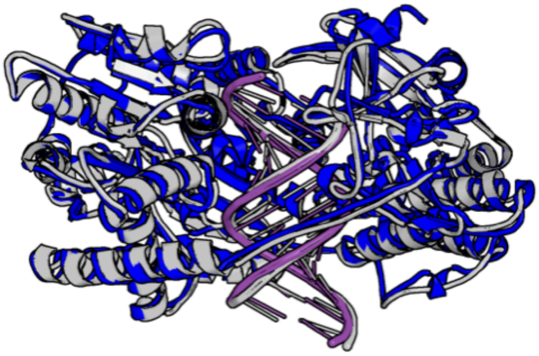

**7TNY (Boltz-1)**
RNA-protein DockQ 0.532/0.592
**RNA-RNA DockQ 0.894**
Total DockQ 0.673
TM-score 0.974

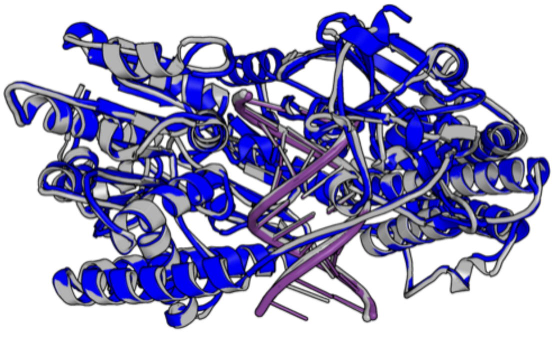

**7TNX (AF3)**
RNA-protein DockQ 0.754/0.805
**RNA-RNA DockQ 0.883**
Total DockQ 0.814
TM-score 0.984

**Figure S10.** Visualization of the top 2 predicted interfaces for both RNA-protein and RNA-RNA interfaces. Top row shows the top 2 predicted RNA-protein interfaces (PDB 8T29 HF3 and 8T29 AF3), while the bottom row shows the top 2 predicted RNA-RNA interfaces (PDB 7TNY Boltz-1 and 7TNX AF3) with IF-DockQ scores shown in bold.

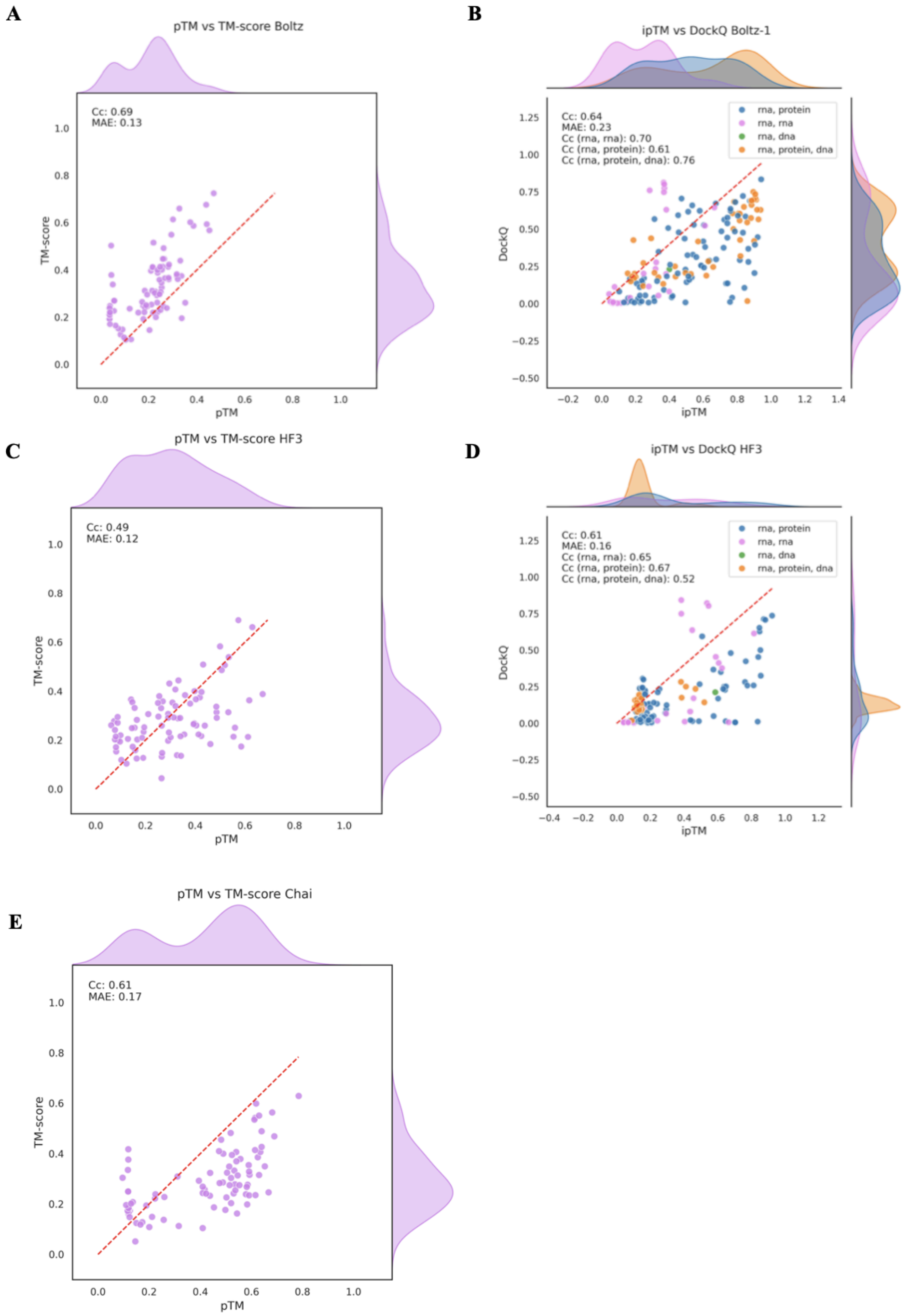


**Figure S11.** Evaluation of pTM/ipTM confidence scores for (A) Boltz and single-chain RNAs, (B) Boltz and RNA complexes, (C) HelixFold3 and single-chain RNAs, (D) HelixFold3 and RNA complexes, (E) Chai and single-chain RNAs. For single-chain RNA correlation between TM-score and pTM is shown, while correlation between IF DockQ and ipTM is shown in the case of RNA complexes. For all methods, RNA-RNA complexes are colored in pink, RNA-protein in blue and RNA-DNA in green.

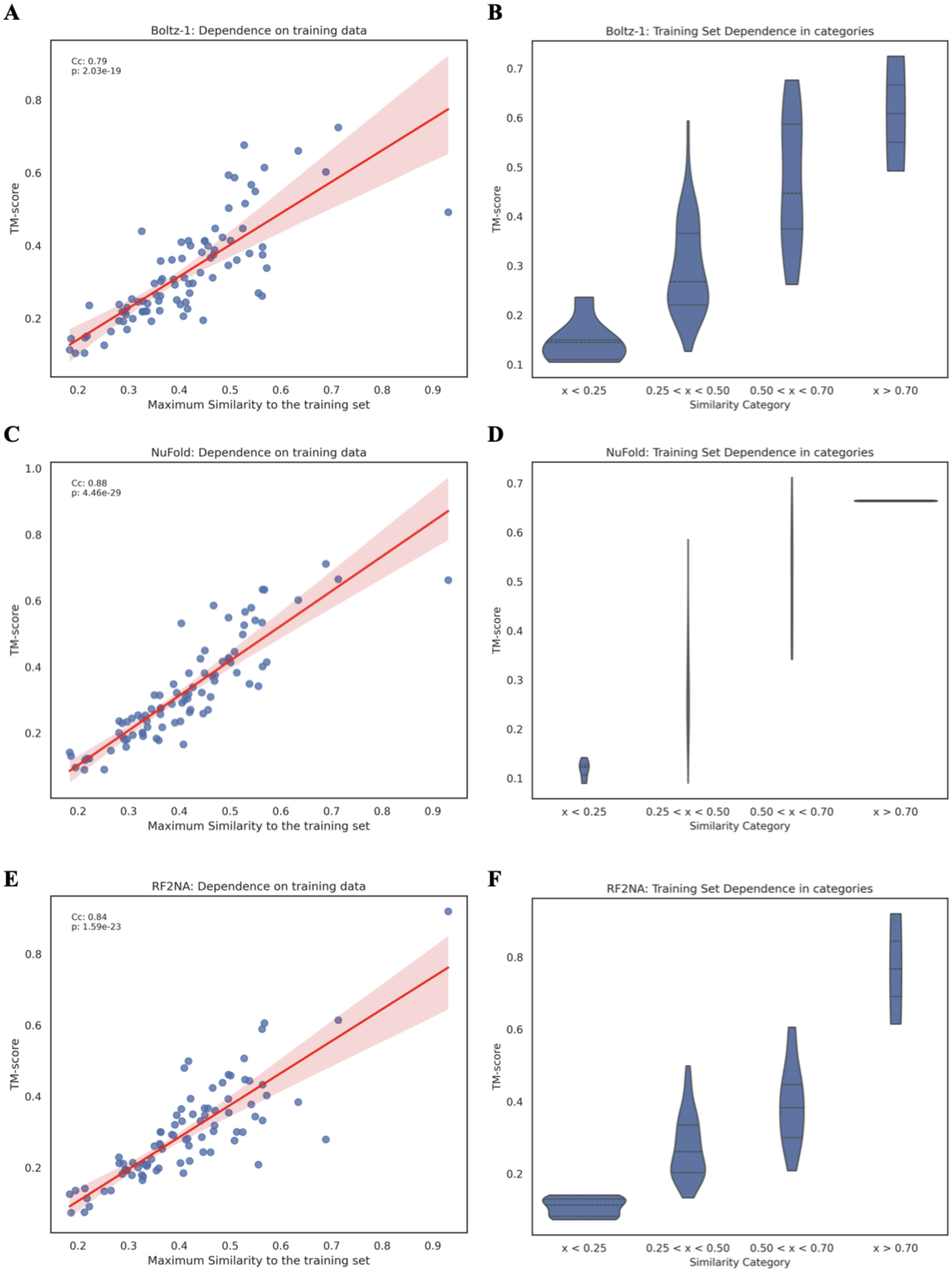


**Figure S12.** The relationship between prediction accuracy (TM-score) and maximum structural similarity to the training set for each single-chain RNA for (A) Boltz-1, (C) NuFold and (E) RF2NA. Prediction accuracy stratified by four training set similarity categories (TM-score: < 0.25, 0.25 - 0.5, 0.5 - 0.75, > 0.70) for (B) Boltz-1, (D) NuFold and (F) RF2NA

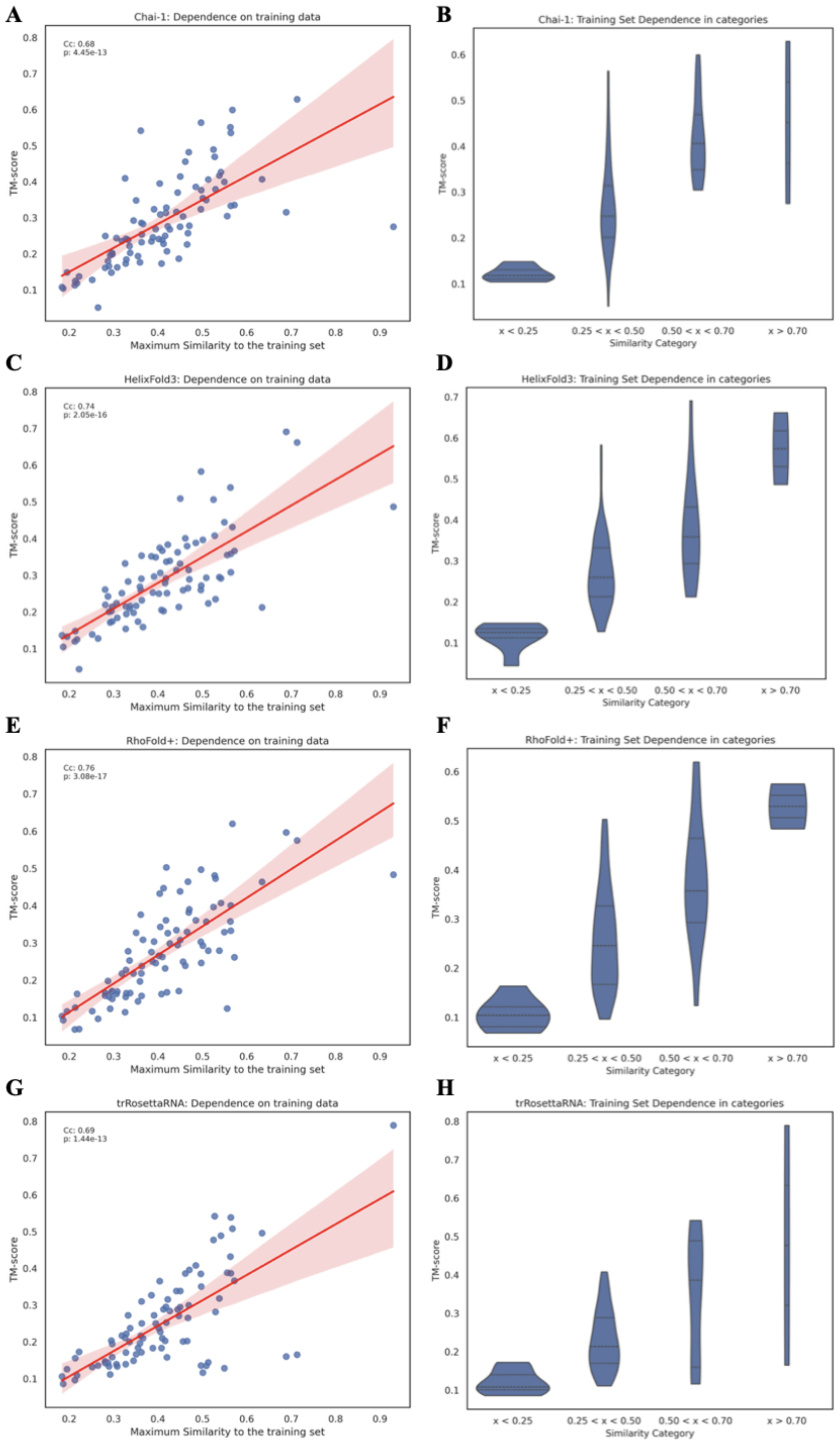


**Figure S13.** The relationship between prediction accuracy (TM-score) and maximum structural similarity to the training set for each single-chain RNA for (A) Chai-1, (C) HF3, (E) RhoFold+ and (G) trRosettaRNA. Prediction accuracy stratified by four training set similarity categories (TM-score: < 0.25, 0.25 - 0.5, 0.5 - 0.75, > 0.70) for (B) Chai-1, (D) HF3, (F) RhoFold+ and (H) trRosettaRNA
